# A regulatory circuit motif dictates whether protein turnover fluxes are more predictive as biomarkers than protein abundances

**DOI:** 10.1101/2021.07.19.452900

**Authors:** Paul M Loriaux, Ying Tang, Alexander Hoffmann

## Abstract

The identification of prognostic biomarkers fuels personalized medicine. Here we tested two underlying, but often overlooked assumptions: 1) measurements at the steady state are sufficient for predicting the response to drug action, and 2) specifically, measurements of molecule abundances are sufficient. It is not clear that these are justified, as 1) the response results from non-linear molecular relationships, and 2) the steady state is defined by both abundance and orthogonal flux information. An experimentally validated mathematical model of the cellular response to the anti-cancer agent TRAIL was our test case. We developed a mathematical representation in which abundances and fluxes (static and kinetic network features) are largely independent, and simulated heterogeneous drug responses. Machine learning revealed predictive power, but that kinetic, not static network features were most informative. Analytical treatment of the underlying network motif identified kinetic buffering as the relevant circuit design principle. Our work suggests that network topology considerations ought to guide biomarker discovery efforts.

**Graphic abstract:** 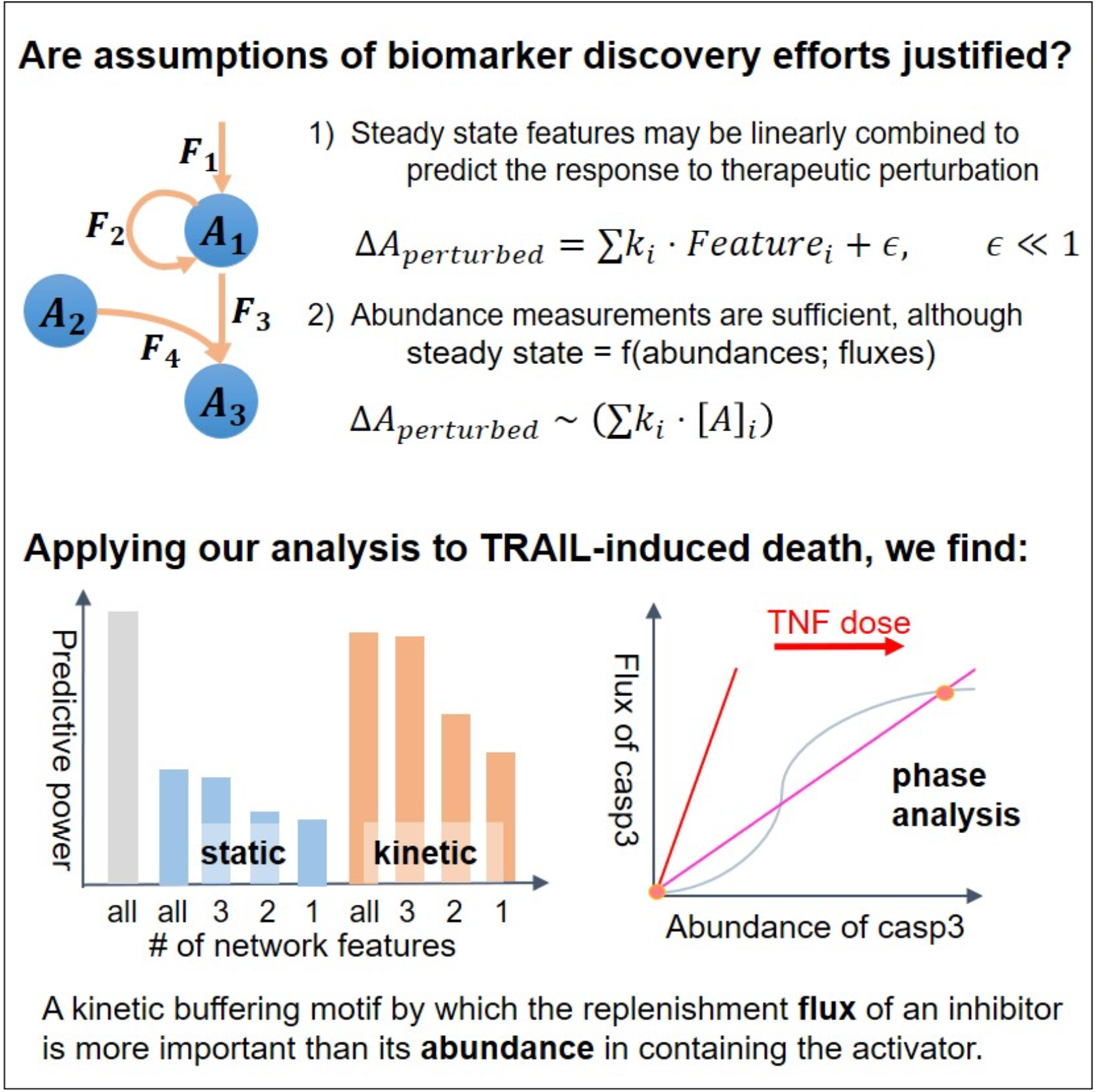

**Highlights:**

– Biomarkers are usually molecule abundances but underlying networks are dynamic
– Our method allows separate consideration of heterogeneous abundances and fluxes
– For the TRAIL cell death network machine learning reveals fluxes as more predictive
– Network motif analyses could render biomarker discovery efforts more productive

**eTOC blurb:** Precision medicine relies on discovering which measurements of the steady state predict therapeutic outcome. Loriaux et al show – using a new analytical approach – that depending on the underlying molecular network, synthesis and degradation fluxes of regulatory molecules may be more predictive than their abundances. This finding reveals a flaw in an implicit but hitherto untested assumption of biomarker discovery efforts and suggests that dynamical systems modeling is useful for directing future clinical studies in precision medicine.

## Introduction

The development of prognostic molecular biomarkers to predict the response to treatments is dramatically improving the diagnosis and treatment of cancer (Auffray et al., 2009; Hood and Friend, 2011), and many other diseases. Distinct from efforts to identify genetic variants that are predictive of disease, biomarker discovery efforts are focused on relating the measured abundances of biomolecules to the efficacy of a specific treatment (Rifai et al., 2006). For example, expression of the receptors for estrogen, progesterone, or epidermal growth factor predict which breast cancers are effectively treated by the adjuvants Trastuzumiab or Tamoxifen (Piccart-Gebhart et al., 2005; Romond et al., 2005), and molecular signatures predictive of treatment efficacy have been identified for histologically indistinguishable B-cell lymphomas (Alizadeh and Staudt, 2000; Rosenwald and Staudt, 2003).

In spite of these successes, the predictive power of such molecular signatures often remains limited (Chechlinska et al., 2010). The common interpretation is that the clinical variability needs to be further stratified into a larger number of distinct classes, prompting further biomarker discovery pipelines (Diamandis, 2010). We wondered whether the focus on abundance measurements in ongoing biomarker discovery efforts was justified or might instead limit their success. After all, even if the steady state of a dynamical system (such as the regulatory network controlling cell proliferation or death) is predictive of its response to a perturbation (such as an anti-cancer drug), fundamental theory states that the steady state is only incompletely determined by the abundances of all of its components; the fluxes and turnover rates provide non-redundant, potentially critical information. Indeed, a recent study demonstrated that turnover rates may vary between cells and samples (Alber et al., 2018). As such, using molecular abundances as biomarkers for therapeutic decision-making could potentially inherently limit predictive power.

To address this question and test the implicit assumption that molecular abundances are sufficiently predictive -we sought a cancer therapeutic whose mode of action was sufficiently well-defined that it was recapitulated in a kinetic mathematical model of the molecular network, so that we could interrogate the afore-mentioned questions in a more systematic and detailed manner than is possible with any experimental model system. The anti-cancer therapeutic, rhTRAIL/APO2L (hereafter, “TRAIL”) is a TNF family member that preferentially induces apoptosis in transformed cells (Wiley et al., 1995). Several TRAIL analogues have been tested clinically (Ashkenazi and Herbst, 2008), but the response to TRAIL is heterogeneous and cell-type specific (Mahalingam et al., 2011; Wagner et al., 2007). An experimentally validated mathematical model of TRAIL-induced cell death based on ordinary differential equations (ODEs) was developed (Albeck et al., 2008a), and extended to include protein synthesis and degradation for all molecular species that produces a non-trivial steady state of live cells that may withstand or die in response to TRAIL treatment (Loriaux et al., 2013).

Here, we developed a computational framework that combines bottom-up network analysis with top-down machine learning to identify a parsimonious set of biochemical features that accurately predict the response to chemical perturbation in a heterogeneous population of cells. To address the specific question about the predictive power of abundance information, we reformulated the kinetic model in terms of abundances and turnover parameters, using a newly developed method called py-substitution (see Box). In contrast to conventional formulations using kinetic parameters which control both the abundance and flux of a molecular species, in our formulation fluxes are controlled by balancing synthesis and degradation rates to insulate them from abundances. Hence, the model formulation distinguishes between static (abundance) and kinetic (flux) features.

By letting protein abundance and flux parameters be sampled from empirically-derived distributions, we were able to rapidly evaluate the response to TRAIL for a diversity of different cellular steady states that reflect the potential diversity of tumors. Using quadratic programming feature selection, we first confirmed that steady state information can indeed serve as an accurate predictor of the response to TRAIL, independent of the stimulus-induced dynamics of biochemical reactions. Remarkably, however, we found that the features with the most predictive power are kinetic rather than static. Indeed, four kinetic features provided 78% prediction accuracy. That is, we show that the rates of synthesis and degradation of just two key regulatory molecules are considerably more informative than the absolute abundances of those or in fact all molecules combined. Further, analytical treatment of a minimal model identified the network motif that renders kinetic features particularly important in the TRAIL death network. Our results have strong implications for clinical prognosis of drug sensitivity; they argue that at least for some clinical presentations – depending on the underlying regulatory network – significant predictive information is lost when live clinical specimens are fixed after biopsy instead of being directly used measurements.

## Results

### Testing assumptions of biomarker discovery efforts using drug-responses to TRAIL as an example

Biomarker discovery efforts are based on the assumption that measurements of specific features of the relevant biological network at steady state may be linearly combined to predict the response to therapeutic intervention, and further, that information of the abundances of network components (necessitated by commonly available measurement tools) is sufficient, although the steady state is of course a function of both abundances and fluxes (Figure 1A). To address these assumptions implicit in biomarker discovery efforts, we developed an analytical workflow that combined (i) high throughput mechanistic mathematical modeling heterogenous patient cancers, (ii) machine learning feature selection by data-driven modeling, and (iii) model reduction to identify design principles using analytical tools (Figure 1B). Key to being able to address the assumption that molecule abundances provide sufficient predictive power is our recently developed method, termed py-substitution (Box), to mathematically transform dynamical systems model written in a mass action formalism in a manner that separates the parameters that control molecule abundances and fluxes (Loriaux and Hoffmann, 2013; Loriaux et al., 2013). We first demonstrated the workflow with a small model of an enzymatic switch that shows bistability (Supplementary Figures 1-2, Methods) before applying it to a larger, published mathematical model that describes the anti-cancer treatment by the biological therapeutic TRAIL (Almasan and Ashkenazi, 2003; Wang and El-Deiry, 2003).

**Figure 1.**
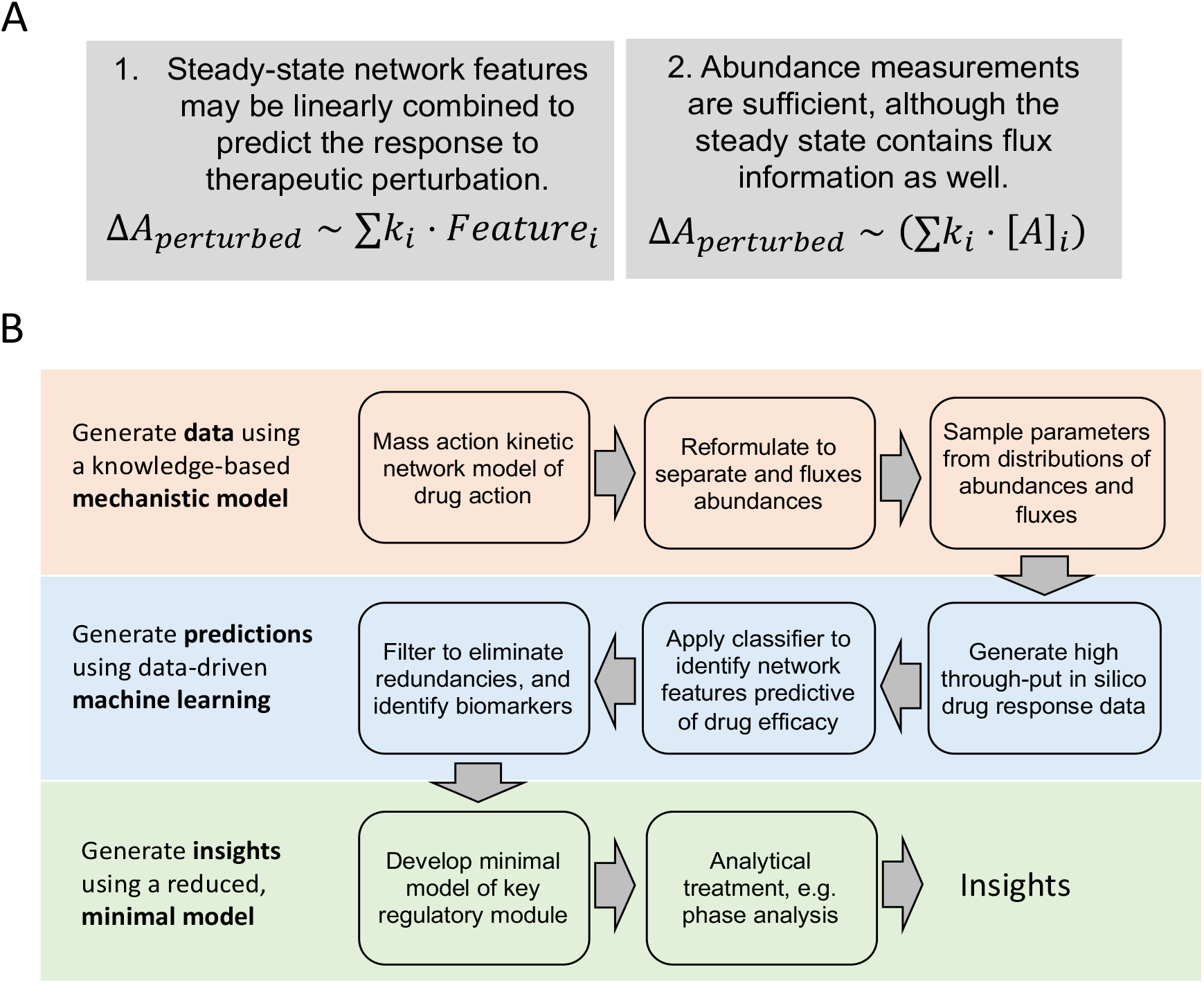
Testing assumptions of biomarker discovery using knowledge-based and data-driven modeling. (A) Two assumptions underlying but often overlooked in biomarker discovery efforts. Biomarker discovery efforts assume that (Left) measurements of the (quasi-)steady state prior to therapeutic intervention may predict the response to treatment although the response is governed by non-linear relationships between molecular species, and (Right) that measurements of molecule abundances is sufficient although the steady state also contains non-redundant, orthogonal information about molecule fluxes or turnover. (B) Schematic of the computational workflow to test the assumptions. An experimentally validated mass action kinetic model based on mechanistic knowledge is reformulated to separate abundance and flux of the steady state as distinct parameters that are then assigned values from distributions for simulations of the heterogeneous responses of a population. The resulting data is applied to data-driven machine learning classification to identify informative, non-redundant features of the steady state that are predictive of drug responses. Based on the identified features a minimal mechanistic model is constructed for analytical treatment to understand why they are so informative.

### Py-substitution – a method to separate static and kinetic network features

Mathematical models of molecular networks typically describe a set of chemical reactions, as rate equations by the Law of Mass Action (Sreenath et al., 2008): 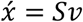, where 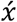 denotes the change in abundance of species over time, *S* is the stoichiometric matrix and *v* is the reaction flux, which is given by the product of kinetic rate constants *k* and the abundance at that time. Thus, in the mass action formulation kinetic rate constants determine the abundances of each molecular species both at steady state as well as in response to therapeutic perturbation. In other words, rate constants are independent parameters and abundances depend on them.

However, measurements of the steady state are typically molecular abundances, especially when biopsy tissue is fixed or frozen for preservation purposes. That means we require a formulation of the mathematical model in which abundances are independent parameters. If abundances are independent, then kinetic information is specified by the flux, as flux may change without altering abundances. Therefore, we seek a formulation of the mathematical model that allows simulating the response to perturbation as a function of abundances and fluxes, rather than kinetic rate constants.

Py-substitution is a method to substitute kinetic rate constants with abundances and fluxes, by deriving an analytical expression for the steady state. The previous method of deriving analytical expressions for the steady state was restricted to the case with flux as a degree-1 function of the abundances (King and Altman, 1956; Lam and Priest, 1972; Volkenstein and Goldstein, 1966). In signaling systems, the flux *v* is often a function of higher degree abundances, as regulation may involve complexes with multiple components and stoichiometries. The py-substitution can derive steady state expressions for models where the flux *v* is degree 1 in reaction rates and degree 2 in abundances. That enables the derivation of analytical expression for the steady state of more mathematical models, particularly those describing regulatory or signaling networks.

As illustrated in the box figure, py-substitution involves the following operations (Loriaux et al., 2013):

1. Choose *d*_*y*_ (*d*_*y*_ > *rank S*) parameters *y* as a set *Y*. The parameters should be linear in flux *v*, and can be specified by the mapping: *ψ*_*p*_(*v*) = *Ty*, where *T* is the *d*_*k*_ × *d*_*y*_ Jacobian matrix with *T*_*ij*_ = *∂v*_*i*_/*∂y*_*j*_. Then, the steady state condition becomes *Cy* = 0, with *C* ≐ *ST*. The remaining parameters are kept identical in a set *P* under the mapping.
2. The second step is to further partition the set *Y* for identifying dependent parameters therein. It is achieved by Gaussian elimination on matrix *C*. For the reduced row echelon form *C*, the columns with a pivot identify the position of dependent parameters in *Y*, and those without a pivot correspond to independent ones. Let matrix *N* have the basis for the null space of *C*, and thus *y* = *Nq* satisfies *Cy* = 0. Then, the mapping through *y* = *Nq* separately maps the set *Y* to *Q* with the independent parameters and another complementary set spanned by the sets *P, Q*: *span*(*P, Q*). The set *P* is kept identical, and they together define the mapping *ψ*_*y*_.
3. After specifying the dependent parameters, we conduct the inverse mapping of the above two: (*ψ*_*p*_ · *ψ*_*y*_) ^−1^. Since they are reversible (Loriaux et al., 2013), we finish the separation on the independent parameters of the subset *P, Q* and the dependent parameters of *Y*.

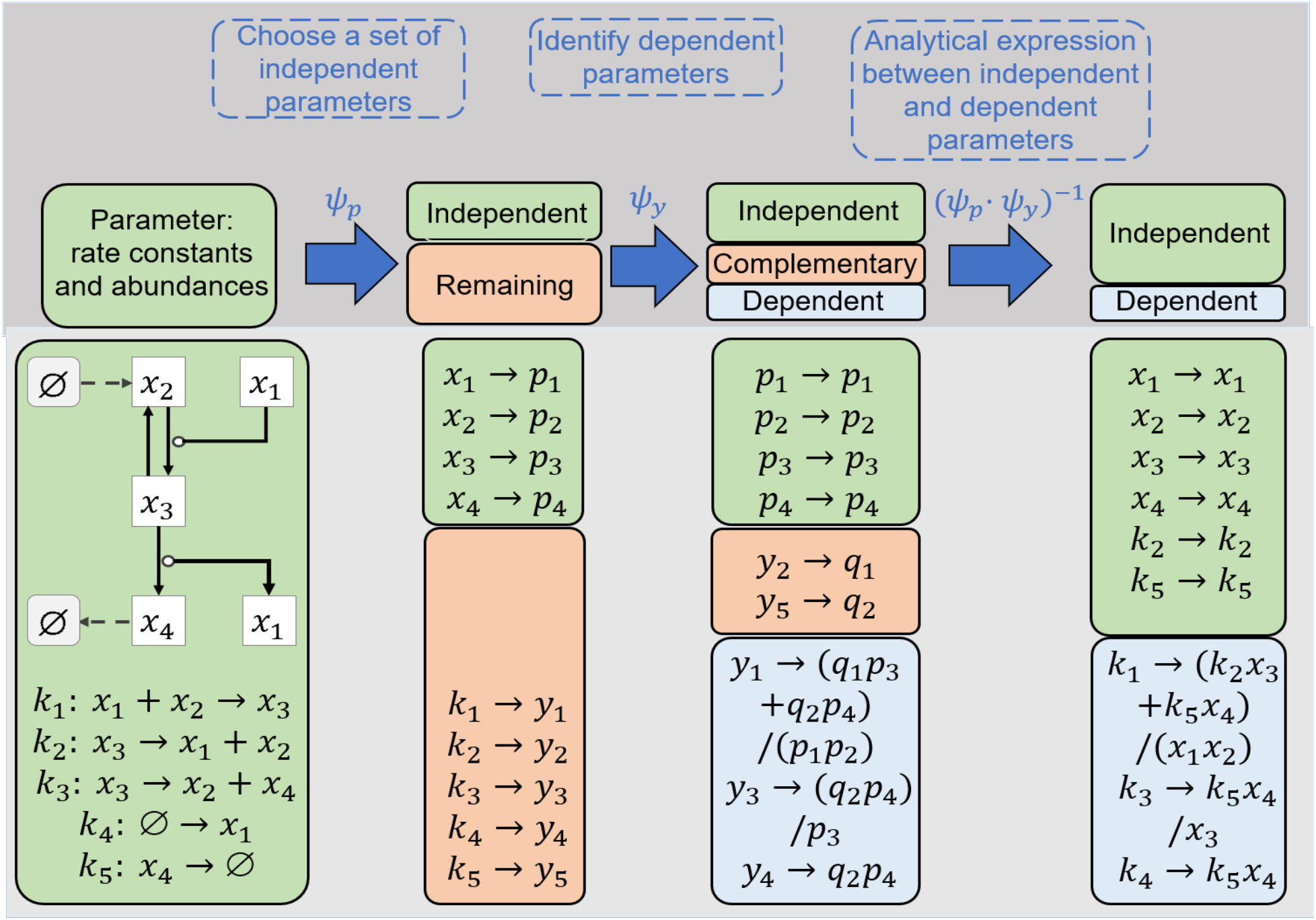

An illustrative example is the open system of Michaelis-Menten enzyme kinetics (Loriaux et al., 2013). The reactions are provided in Supplementary Figure 3, with the rate constants {*k*_*i*_}_*i*=1,…,5_. We can choose the four abundances as the independent parameters in the set *P* = {*p*_*i*_}_*i* =1,…,4_, and the rate constants as the set *Y* = {*y*_*i*_}_*i* =1,…,5_ under the mapping *ψ*_*p*_. The second mapping *ψ*_*y*_ further identifies the dependent parameters in the set *Y*, and here the complementary set in *Y* contains {*y*_2_, *y*_5_}, which are mapped to the set *Q* = {*q*_1_, *q*_2_}. The set *P* is identical under *ψ*_*p*_, and then the parameters in the sets *P, Q* can represent the dependent parameters in the set *Y*. Finally, by conducting the inverse map (*ψ*_*p*_ · *ψ*_*y*_) ^−1^, we obtain the analytical relation between parameters. With the relation, the perturbations on the independent parameters transmit to the dependent parameters, such that the steady state maintains, but not to other independent parameters.

Whereas in conventional parameter sensitivity analysis, changes in kinetic rate constants affect both steady state abundances and fluxes, following py-substitution changes in the independent kinetic parameters generally lead to changes in flux but independent abundances remain unchanged; conversely, changes in abundances will not lead to changes in flux (Loriaux and Hoffmann, 2013). Thus, we are able to study separately the roles of “kinetic features” and “static features” in determining the response to perturbation.

The apoptosis mathematical model consists of ordinary differential equations that describe the time-dependent behavior of 58 molecular species in response to stimulation by TRAIL (Albeck et al., 2008a), extended to include synthesis and decay of every molecular species to produce non-trivial steady states (Loriaux et al., 2013) (Supplementary Figure 3A). Briefly, the model describes TRAIL-mediated assembly of the death-inducing signaling complex (DISC) and subsequent activation of caspase 8. Active caspase 8 cleaves and activates caspase 3, but also Bid, which in turn activates cytoplasmic Bax. Activated Bax then enters the mitochondria to form tetrameric pores through which cytochrome C is released into the cytoplasm. This event is commonly called mitochondrial outer membrane permeabilization, or MOMP. Once in the cytoplasm, cytochrome C catalyzes the formation of the Apoptosome, which results in degradation of XIAP and feed-forward activation of caspase 3. Activation of caspase 3 is required for chromatin condensation and DNA fragmentation (not modeled), and ultimately results in cell death (Porter and Jänicke, 1999; Woo et al., 1998). Because MOMP is irreversible (Sheridan and Martin, 2008), we used the abundances of tetrameric Bax (Bax4) and caspase 3-cleaved poly (ADP-ribose) polymerase (cPARP) as indicators of cell fate (Albeck et al., 2008b; Kaufmann et al., 1993). See Supplementary Tables 1-3 for all species, reactions, and differential equations used in our model.

In order to treat the steady state flux as an independent variable, we used py-substitution (Box) to derive an analytical expression connecting the kinetic rates and abundances at steady state (Loriaux and Hoffmann, 2013; Loriaux et al., 2013). The resulting expression has 119 independent parameters, 18 of which are species abundances, 100 of which are kinetic rate constants, and 1 of which describes the mitochondrial volume. The remaining 40 abundances and 15 rate constants are rational polynomials in the independent parameters by py-substitution. (See Supplementary Tables 4-7 for all parameter values used to numerically integrate the differential equations.) To verify that our model was capable of distinguishing between high and low doses of TRAIL, we generated dose-response curves over a 100-fold increase in ligand abundance. At each dose we simulated the model to 48 hours and recorded the abundance of eight species both receptor proximal and distal (Supplementary Figure 3B-I). The results clearly show two distinct dose-response regimes. At low doses there is a linear response in the DISC, but no caspase activation, cytochrome C release, nor cleavage of PARP. At high doses there is complete cytochrome C release, activation of caspases 3 and 8, and accumulation of cPARP. The model can therefore distinguish between perturbations that do and do not result in cell death.

Next, we modeled the response for distinct steady states, as they might occur in different tumors from different patients. Recent single-cell experiments suggest that protein abundances are gamma distributed (Taniguchi et al., 2010). This distribution arises naturally from Poisson production of messenger RNA, followed by exponentially-distributed bursts of protein translation (Friedman et al., 2006). A remarkable conclusion from this work is that for highly expressed proteins, the variance is proportional to the square of the mean. Using the abundances given in (Albeck et al., 2008a), we calculated the variance, shape, and scale parameters of 14 independent species in our model (Figure 2A-B). By virtue of being constrained to the steady state, 40 dependent species also assumed a probability distribution, albeit with higher variance (Figure 2C). In addition, protein half-lives in murine 3T3 cells are known to be log-normally distributed (Schwanhäusser et al., 2011) (Supplementary Figure 4). We therefore let 11 degradation rate constants follow a log-normal distribution with a coefficient of variation (CV) equal to 0.368, equivalent to a variance of 1 hour in the log-normal distribution of protein half-lives (Figure 2D-E), given that signaling proteins tend to be short-lived (Loriaux and Hoffmann, 2013). Again, by virtue of the steady state constraint, 15 synthesis rate constants assumed a probability distribution as well (Figure 2F).

**Figure 2.**
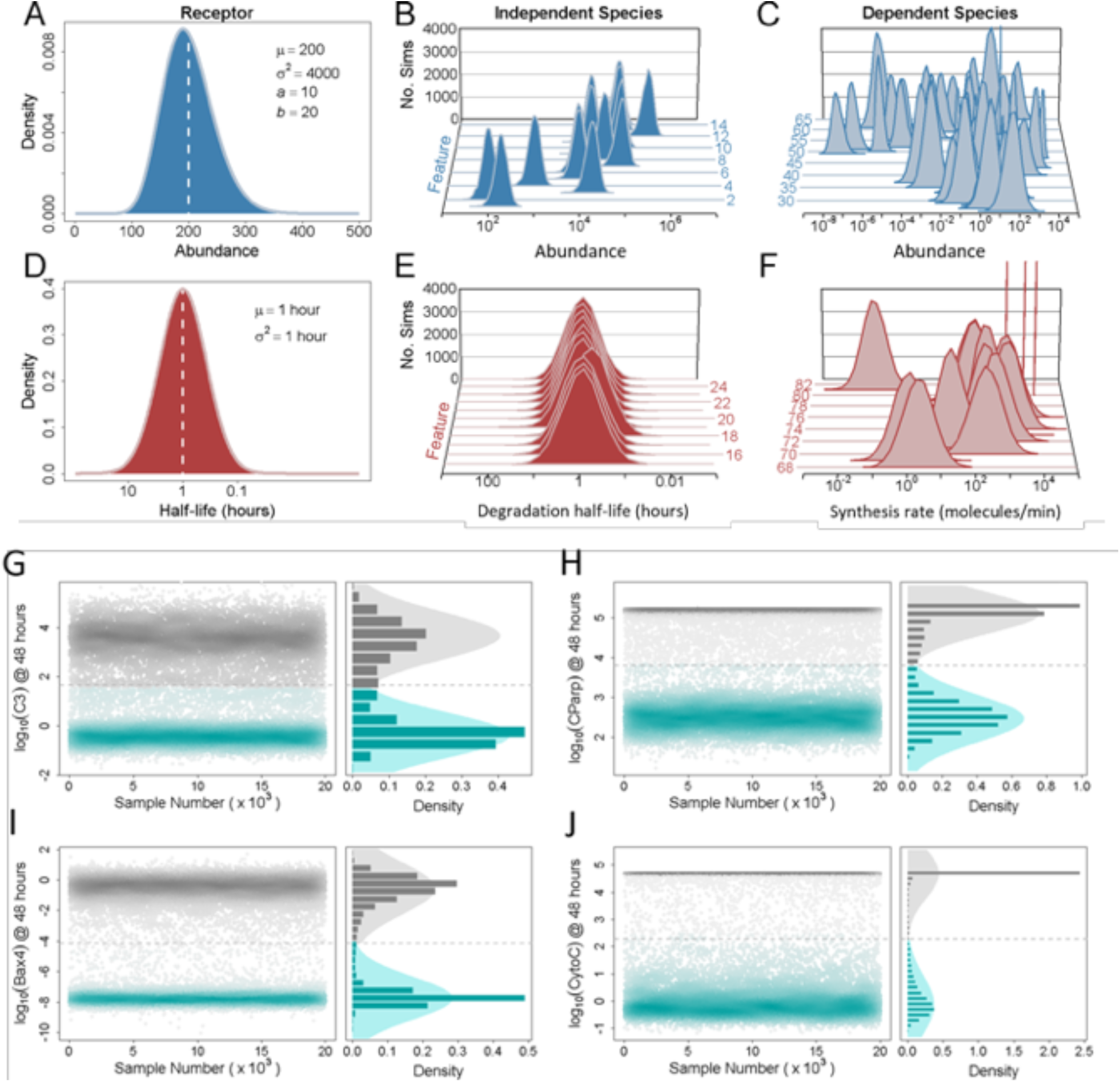
Modeling cell-to-cell heterogeneity with the bistable model of TRAIL-induced cell death. (A) The *a priori* probability density function used for the abundance of the TRAIL receptor. Like other independent species abundances, we let this function be defined by a gamma distribution. The mean *μ* of the distribution is taken to be the value given in (Albeck et al., 2008a); its variance *σ* ^2^ is taken to be one-tenth the square of the mean, i.e. the extrinsic noise limit described in (Taniguchi et al., 2010). From these, a shape *a* and scale *b* parameter were calculated according to the definition of the gamma distribution. (B) *A priori* probability density functions for all 14 independent species abundances. Species are plotted front to back according to their index, as given in Supplementary Table 7. (C) *A posteriori* probability density functions for all 40 dependent species abundances. Each dependent species abundance is a rational polynomial in the independent parameters. (D) *A priori* probability density function for the half-life of the TRAIL receptor. This and other protein half-lives were assigned a standard log-normal distribution with a nominal half-life of one hour, consistent with the observation that signaling proteins are generally short-lived (Loriaux and Hoffmann, 2013). (E) *A priori* probability density functions for all 11 primary protein half-lives, plotted according to their feature index. (F) *A posteriori* probability density functions for the kinetic rate constants describing the efflux of 15 modified proteins and protein complexes. See Supplementary Table 7 for a complete list of all random-valued parameters. Shown in (G-J) are the simulated abundances of four species at 48 hours following the addition of TRAIL, for each of the 20,000 samples. The four species are (G) active caspase 3, (H) cleaved Parp, (I) tetrameric Bax, and (J) cytoplasmic Cytochrome C. For each panel, the left plot shows the absolute abundance of each species as a function of sequential sample number. Points are shaded by density and assigned a color post-analytically based on whether that abundance corresponds to a positive response (gray; cell dies), or a negative response (cyan; cell lives). The right plot in each panel shows a 1-D histogram of the response as well as the pair of Gaussians fitted to its density estimate (see Methods). The saddle point between the two Gaussians is indicated by the dashed line and distinguishes responsive from unresponsive samples.

Next, we sampled the model 20,000 times and simulated its response to an ambiguous dose of TRAIL (1,000 ligands per cell). Examining the abundance of four species --active caspase 3 (Figure 2G), cPARP (Figure 2H), tetrameric Bax (Bax4) (Figure 2I), and cytoplasmic cytochrome C (Figure 2J)--at 48 hours after stimulation revealed two distinct subpopulations. Cells that experienced MOMP achieved the hyperactive steady state indicative of cell death. Cells that did not experience MOMP returned to their pre-stimulated steady states. Due to the symmetry and distance of the two subpopulations of Bax4 at 48 hours, we chose this as our primary response variable. We note that the abundance of Bax4 in the responding population is between 1 and 10 tetramers, in good agreement with the observation that only a few mitochondrial pores are required for MOMP (Düssmann et al., 2010). After fitting with a two-component Gaussian mixture model, those that returned to their pre-stimulated steady states were scored as “alive” and assigned a response value of zero. Those that did not were scored “dead” and assigned a response value of one.

### Steady state can be an accurate predictor of the response to TRAIL

After binarizing the response, we returned to the prior empirical parameter distributions and calculated their correlation with the response. We found that we can build an informative classifier for a heterogeneous population of cells using only ∼10^3^ simulations (Supplementary Figure 5). Note that random-valued parameters in the forward numerical integration problem are considered “features” in the backward model selection problem. We thus distinguish between “static” and “kinetic features”, where static features are the steady state abundances of the molecular species and kinetic features are the synthesis (and degradation) fluxes that when balanced do not affect their steady state abundances. As shown in Figure 3A, no single feature correlates well with the response to TRAIL. This is in agreement with the conclusion drawn by (Spencer et al., 2009), where no single protein exhibits strong correlation with the time elapsed between administration of TRAIL and MOMP, unless artificially overexpressed.

**Figure 3.**
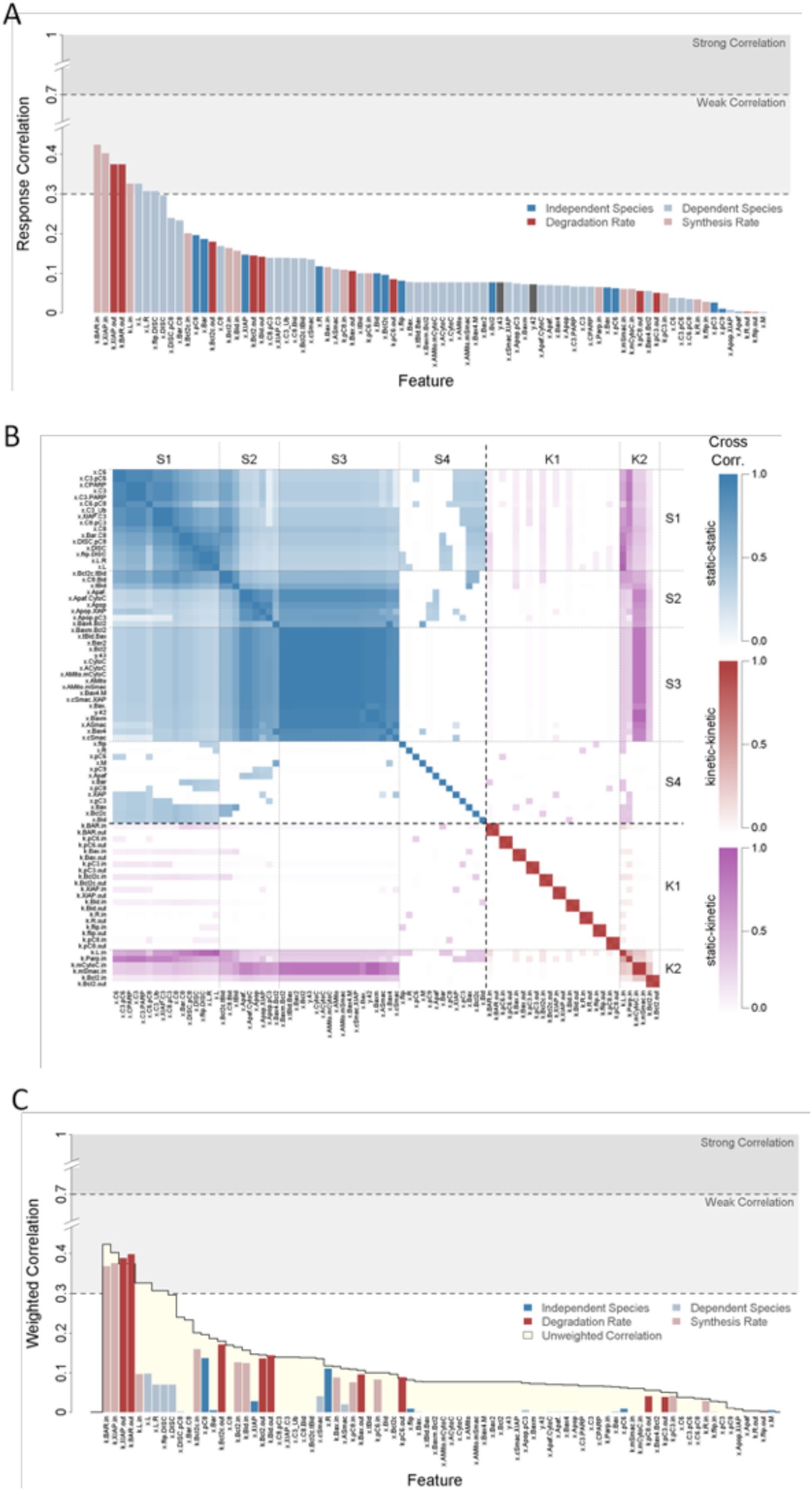
Correlation between features and the binary response. (A) Pearson correlation coefficients were calculated between each feature and the binary response variable derived from the abundance of tetrameric Bax at 48 hours (see Methods). The absolute values of the coefficients are shown here, sorted in decreasing order and color-coded to indicate whether the feature is an independent species (dark blue) or degradation rate constant (dark red), or a dependent species (light blue) or rate of synthesis (light red). Light and dark gray rectangular regions in the plot illustrate regions of weak and strong correlation, respectively. (B) Extensive correlations between features indicate redundancies. For each of the number of simulations given, the correlations between all 82 features and the response variable were calculated and subtracted from the final correlation shown in (A). These were then plotted as boxes. Values close to zero indicate correlations equal or nearly equal to those shown in (A). (C) QPFS weighted correlation between features and the binary response to TRAIL. Features are plotted in the same order as in (A), but bar height now represents the QPFS-weighted correlation between each feature and the response to TRAIL rather than the raw, unweighted correlation. For comparison, the unweighted correlations are indicated here by the yellow silhouetted region. As before, features are color-coded to indicate whether they are independent species (dark blue), dependent species (light blue), degradation rate constants (dark red), or synthesis rates (light red).

A confounding issue of biological systems is that the feature set or measurements are highly cross-correlated. To account for redundancy, we calculated the cross-correlation matrix (Figure 3B). As expected, every feature perfectly cross-correlates with itself, as illustrated along the diagonal. Because steady state abundances and fluxes were separated, paired synthesis and degradation rates are highly correlated as well (clusters K1 and K2). Interestingly, cluster K2 contains fluxes that are well-correlated with a number of species abundances. Among the static features, we observed four clusters. Cluster S4 contains the independent species abundances, all of which exhibit low correlation with every other feature. Clusters S1, S2, and S3 are the receptor-proximal, post-mitochondrial, and mitochondrial clusters, respectively. The high degree of cross-correlation observed in cluster S3 argues that if the state of the mitochondria is a good predictor of TRAIL sensitivity, then only one mitochondrial feature will need to be measured. Similar arguments can be made for clusters S1 and S2. However, molecular species that are well separated in the biochemical network, e.g. activated caspase 6 and cytoplasmic Bcl-2, are nevertheless strongly correlated due to the constraints imposed by steady state.

Using the correlation between each feature with the response, and the cross-correlation between every pair of features, we sought to identify a subset of maximally predictive, minimally redundant features for predicting the response to TRAIL. To do this we used quadratic programming feature selection (Rodriguez-Lujan et al., 2010), or QPFS. QPFS expresses this objective as a quadratic program and finds a vector of weights on the features that minimizes redundancy while maximizing relevance (see Methods). If we examine the QPFS-weighted correlations between each feature and the response (Figure 3C), we see a dramatic reduction in complexity. In particular, above the weighted correlation thresholds of 0.3 and 0.12, we observe only four and twelve features, respectively. These are the rates of synthesis and degradation of BAR and XIAP, and above 0.12 the rates of synthesis and degradation of cytoplasmic Bcl-2, mitochondrial Bcl-2, Bid, and the steady state abundances of procaspase 8 and the TRAIL receptor.

We therefore asked whether we could accurately predict the response to TRAIL using only the small subsets of features. We used logistic regression to model the log-odds ratio of the probability of responding to TRAIL as a linear combination of the four or twelve features. To do this we divided the 20,000 simulated responses into equal sized training and test datasets. Regression coefficients were derived by maximum likelihood estimation on the training data. We then compared predictions from the logistic regression models with the test data and found that they achieved 78 and 84 percent accuracy, respectively (Figure 4A-B). This indicates that, when predicting the response to TRAIL, a substantial reduction in model complexity (linear combination of a small set of features rather than 54 non-linearly related species) can be achieved with only a modest loss in accuracy. In sum, the first assumption of biomarker discovery efforts (Figure 1A) appears to hold true for this example of TRAIL-induced death.

**Figure 4.**
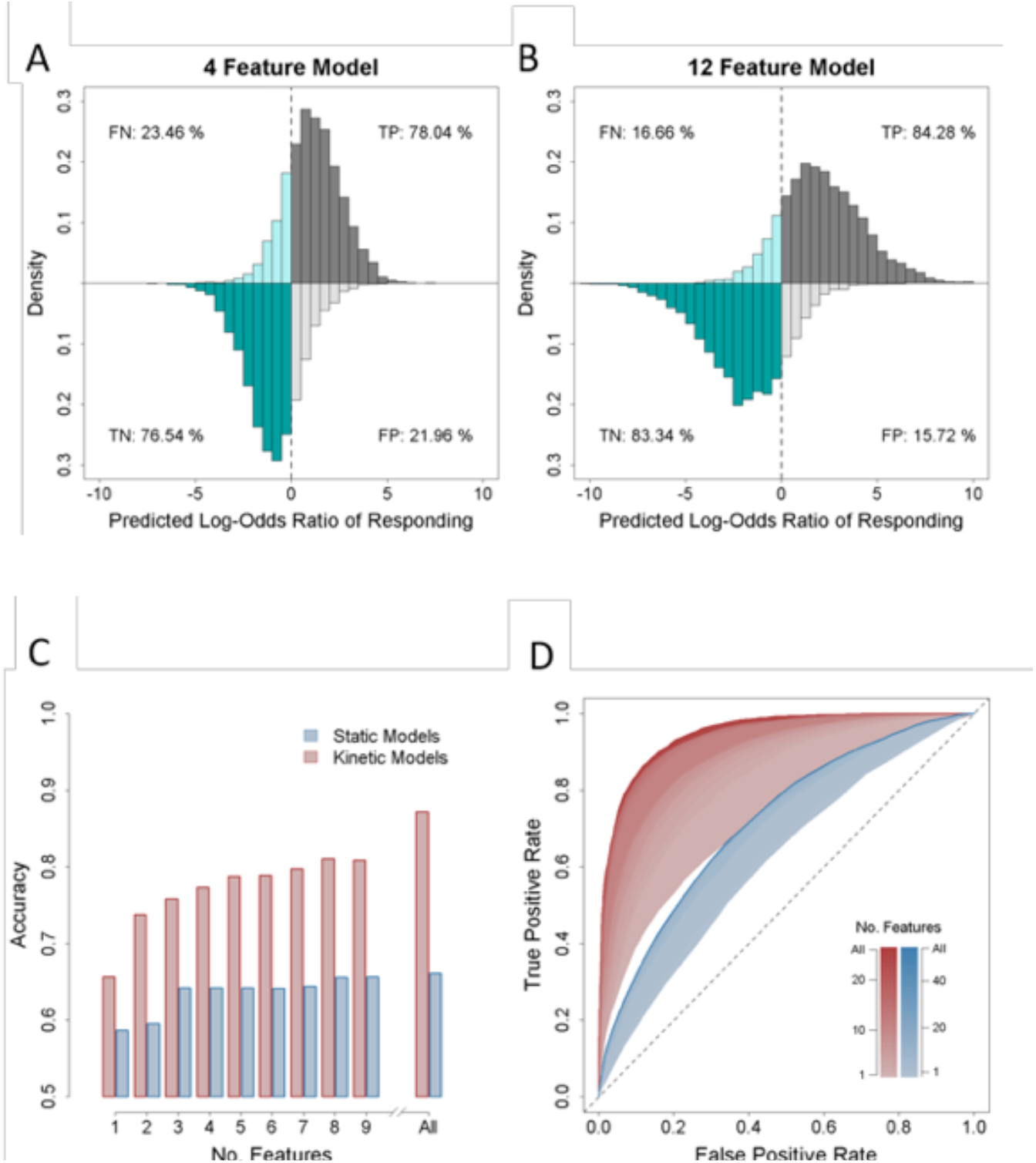
Kinetic features outperform static features. (A) The four top-ranked features as identified by QPFS were used to build a logistic regression model. In decreasing order, those features are the rates of synthesis of BAR and XIAP, and the rates of degradation of XIAP and BAR. The model was trained on one half of the dataset of 20,000 simulations and tested on the other. The abscissa gives the predicted log-odds ratio of the probability of responding to TRAIL versus not responding. A value of zero corresponds to equal probability. Values greater than zero are predicted to respond to TRAIL (cells die; gray), while values less than zero are predicted not to respond (cells live; cyan). Samples in the first and second quadrant are those that actually responded to TRAIL according to the ODE model (positives). Samples in the third and fourth quadrant are those did not respond to TRAIL (negatives). (B) As above, but the twelve top-ranked features as identified by QPFS were used instead. These include: the rates of synthesis and degradation of cytoplasmic Bcl-2, mitochondrial Bcl-2, Bid, and the steady state abundances of procaspase 8 and the TRAIL receptor. (C) Static-only (blue) or kinetic-only (red) features were iteratively added to a logistic regression model in order of their QPFS ranking and the models trained and tested as before. The accuracy of each model as a function of its size is shown here, where “All” indicates either all 56 static features or all 26 kinetic features. An accuracy of 0.5 is equivalent to random guessing. (D) For each of the 56 static and 26 kinetic regression models, a receiver operating characteristic curve was generated by varying the classification threshold for responders and non-responders over the entirety of its range.

### Kinetic features outperform static features as response predictors

Interestingly, our results also suggest that kinetic features are more powerful predictors of TRAIL-responsiveness than static features. All seven of the best-performing features are kinetic. Since only 33% (27 out of 82) of all features are kinetic, this enrichment is remarkable. This result is largely robust to when turnover rates are substantially slowed (Supplementary Figure 6). To further examine the predictive power of kinetic versus static features, we built all 26 and all 54 logistic regression models using only kinetic or static features, respectively, in decreasing order of their QPFS ranking. We then calculated the accuracies of these models as before (Figure 4C). The results confirm that kinetic regression models outperform static regression models. The regression model built using only the rates of synthesis and degradation of XIAP is superior in accuracy to the model built using all 54 steady state molecule abundances.

An implicit parameter in the calculation of model accuracy is the threshold at which an unknown system is classified as responsive. By convention this threshold is zero, that being when the predicted probability of responding equals the probability of not responding. By altering this threshold, the fraction of true to false positives can be adjusted, yielding the well-known receiver operating characteristic (ROC) curve. To test whether kinetic regression models simply outperform static models at a particular classification threshold, we generated ROC curves for all 80 logistic regression models. Again, the results confirm that kinetic regression models are far more discriminatory than static regression models. As with the previous result, a kinetic regression model containing only the rates of synthesis and degradation of XIAP outperforms the static regression model incorporating all 54 steady state species abundances (Figure 4D). In sum, the second assumption implicit in biomarker discovery efforts (Figure 1A) appears not to be justified in this example of TRAIL-induced death.

### A minimal model identifies a network motif controlled by kinetic features

To examine why the outcomes of the TRAIL death decision network were more predictably determined by kinetic than static features, we developed a simplified model of the key regulatory step. Besides the apoptosis model considered here (Loriaux et al., 2013), there are other formulations adapted to match protein measurements in different cell types (Albeck et al., 2008a; Lopez et al., 2013; Spencer et al., 2009). A common core module in determining the apoptosis decision is the C3* protein and the regulation between XIAP and C3*. Thus, we considered the interaction between XIAP and C3*, as the C3* response determines the decision on live and death, through catalyzing the reaction from PARP to cPARP (Figure 5A). Activated caspase 3 is modeled within a positive feedback loop which also abstracts positive mitochondrial feedforward loop emanating from caspase 8 in the extended cell death model (Albeck et al., 2008a), and is inhibited by XIAP that can be synthesized and degraded (Loriaux et al., 2013; Spencer et al., 2009). The model is simulated by applying an activation function to C3* which is the result of TRAIL interacting with its receptor, activating caspase 8 and then caspase 3. As such different doses of TRAIL lead to different C3* activation amplitude (Figure 5B, top) that result in reductions of XIAP (Figure 5B, middle). In our example, the highest activation function results in an irreversible depletion of XIAP and activation of C3* (Figure 5B, bottom), signifying death.

**Figure 5.**
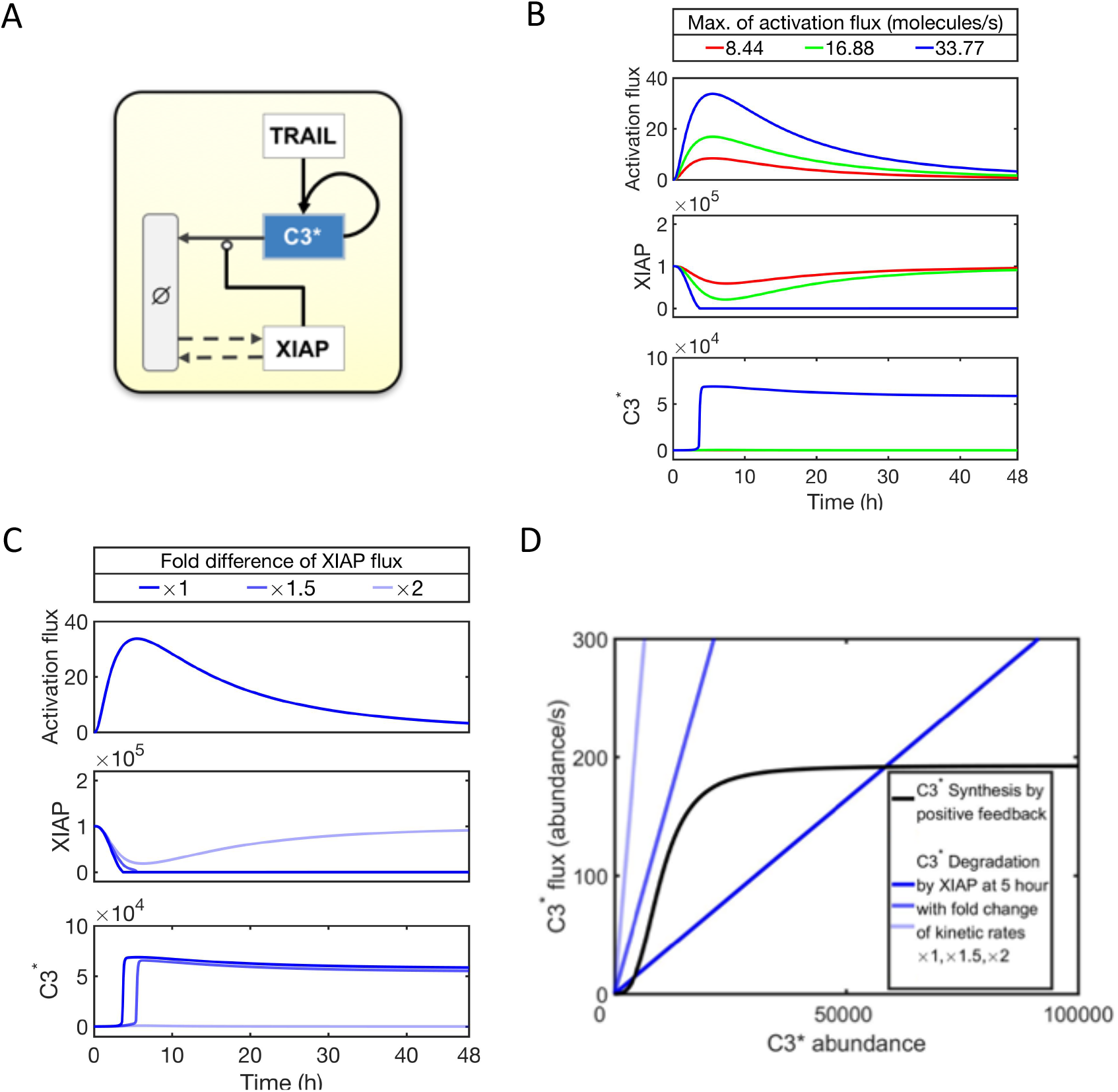
Minimum model provides a mechanistical understanding on the role of kinetic features. (A) The network structure of a minimal model extracted from the detailed cell death model (Supplementary Figure 3A). TRAIL’s effect on C3* is approximated by a pulse-like activation profile. A positive feed-forward loop is present in the whole network (Albeck et al., 2008a) and coarse grained here. (B) Time course of the input activation flux induced by TRAIL, and the corresponding time course of XIAP and C3* abundance. The colors denote fold-change of the activation flux. (C) Time course of XIAP and C3* abundances for a fixed input activation flux by TRAIL, under various of XIAP synthesis and degradation rates. The brightness of lines denotes fold-change of XIAP synthesis and degradation rates simultaneously. For each condition, we used py-substitution to ensure the initial abundances are always at steady state. (D) The plot with synthesis and degradation flux lines of C3* showing its dynamical stability structure. C3* is activated by pulse-like input, which triggers the synthesis flux by positive feed-forward loop. C3* is consumed by the reaction XIAP. If XIAP synthesis and degradation rates are high (low), XIAP concentration is reduced slower (faster), leading to more (less) degradation on C3*. Once the total consumption of C3* is sufficiently low, the positive feed-forward loop dominates, pushing the system reach the high C3* abundance steady state denoting cell death.

We probed the minimal model to understand why kinetic features outperformed the static abundances in determining the binary decision for cell response. When XIAP synthesis and degradation rates are increased at this previously lethal dose of TRAIL activation, XIAP is effectively replenished when an increase of C3* consumes it (Figure 5C). Specifically, increasing XIAP turnover is able to slow the response to the increased C3* activity and when XIAP turnover is increased further, XIAP never reaches the point of complete diminishment, thereby keeping the C3* positive feedback loop in check. As the TRAIL-induce activation function decays after some time, XIAP recovers to the original high abundance and C3* never rises to lethal levels (Figure 5C). Thus, increased turnover of XIAP, without changing its steady state abundance, results in increased resistance to TRAIL triggered pro-apoptotic signaling.

We plotted the flux lines of C3* due to its TRAIL-induced synthesis and inhibition by XIAP (Figure 5D and Supplementary Video). The positive feedback leads to a sigmoidal flux curve and thus an ultra-sensitive relationship of C3* flux vs C3* abundance. Together with the degradation flux caused by XIAP, they define the two regimes of the binary fate decision. With a lethal amount of TRAIL activation, C3* levels drop below the threshold unless its degradation by XIAP is decreased. While a 1.5 x increase does not quite replenish C3* levels entirely, a two-fold increase does, confirming the sensitivity of this kinetic feature of the network. Indeed, the converse can be shown similarly. At TRAIL activation doses that were not lethal (Figure 5B), reducing the turnover of XIAP can render cells substantially more prone to apoptosis. At the sublethal activation dose, a halving of the the flux resulted in irreversibly diminished XIAP and elevated C3* levels (Supplementary Figure 7A); when the activation dose was further reduced, halving the turnover rate was insufficient, but diminishing it by 10-fold resulted in death (Supplementary Figure 7B). Together, we observe that the flux of XIAP is a strong determinant of cell death decisions, as it controls engagement of the C3* positive feedback loop. In contrast, the abundance of XIAP does not alter the cell fate decisions for either lethal or non-lethal activation doses (Supplementary Figure 7C-D). However, when the XIAP inhibitor functions enzymatically rather than stoichiometrically, its steady state abundance is a more potent determinant of cell death than its flux (Supplementary Figure 7E-F).

## Discussion

Here we have examined two common assumptions implicit in biomarker discovery efforts: that a regression model of measurements, in essence a linear representation of the dynamical systems network, retains predictive power about the response to treatment to guide clinical decision making; and further, that measurements of molecular abundances are sufficient to achieve this predictive power, even in the absence of kinetic information. We addressed these assumptions for a potential anti-cancer treatment that is based on the cell-death-inducing ligand TRAIL, and found the answers to be a qualified “Yes” to the first, and a clear “No” to the second. In other words, while linear regression modeling could retain predictive power, the absence of kinetic information about the network substantially limited its performance. In fact, a single turnover or flux measurement could provide greater accuracy and precision than abundance information of all molecules represented in the network.

A quantitative evaluation of the performance of a biomarker requires that the “ground-truth” is known. This can only be provided by a mathematical model of a biological network, not the actual biological specimens, which can provide error-free “measurements” of the biochemical state of the network and the outcome of the drug perturbation. We developed a computational workflow for identifying a parsimonious set of specific biochemical features (i.e. abundances and fluxes) that optimally predicts the response to the drug perturbation within a heterogeneous population. This workflow was made possible by the following recent advances: 1) an experimentally validated reaction network model of the biological process in question (Albeck et al., 2008a), 2) computational methods for deriving an analytical expression for the steady state of the network that allows us to separate static vs kinetic features of the network (i.e. molecular abundances and flux/turnover rates, respectively) (Loriaux et al., 2013), 3) incorporating kinetic variability into modeling (Loriaux and Hoffmann, 2013), and 4) a method to identify subsets of features that are simultaneously predictive and non-redundant (Rodriguez-Lujan et al., 2010).

After applying our workflow to the anti-cancer drug TRAIL, our first conclusion is that information about the steady-state of a dynamical system can indeed serve as a good predictor of its response to perturbation (Figure 3A and B). That is, a logistic regression model that considers only four steady state features will accurately predict, four times out of five, the response to TRAIL compared to the full ODE model. *A priori*, there is no reason to think that this should be possible. It will be of interest to examine whether other biological networks are less well represented by linear regression models. Given that to date, over 500 curated, quantitative models of cell decision and signaling processes have been developed (Le Novère et al., 2006), the potential impact of interrogating them with the described workflow could be substantial.

For TRAIL, we find that the rates of protein synthesis and degradation, or flux, of XIAP and Bar are the strongest predictors of the response. These are followed by the fluxes of Bid and Bcl-2 and the steady state abundances of the TRAIL receptor and procaspase 8. The locality of these features in the reaction network lends credibility to this finding. Bar and XIAP are at the points of bifurcation and convergence of the mitochondrial feed forward loop, respectively. The original ODE model was trained on data derived from HeLa cells, for which MOMP is normally required for cell death (Mandal et al., 1996). The turnover of proteins that control this bifurcation are therefore good predictors of the response.

We conclude that kinetic features of the regulatory network are stronger predictors of TRAIL-induced cell death than static features. This conclusion does not appear to be sensitive to the mean of the kinetic feature distribution (Supplementary Figure 6). Whether the global protein-half life is modeled at 1 or 66 hours, QPFS identifies the fluxes of Bar and XIAP, or Bax and Bcl-2 respectively, as the most powerful predictors of TRAIL responsiveness. In contrast, when the CV of the kinetic feature distribution is reduced from 0.386 to 0.25, as reported in (Gaudet et al., 2012), we see a slight reduction in the dominance of kinetic features. This observation is not surprising, however. In the limit where the variance of a feature is so small as to be effectively constant, knowing that feature will have less impact on classification. Since genome-wide measurements of protein half-life suggest a CV between 0.48 and 2.28 (Boisvert et al., 2012; Cambridge et al., 2011; Schwanhäusser et al., 2011), we conclude that, in general, kinetic features dominate static features when predicting the response to TRAIL. Indeed, static features that were treated as independent random variables in the present study have been shown to be cross-correlated (Gaudet et al., 2012). Incorporating these empirical cross-correlations into the QPFS rankings would further dilute their predictive power and contribute to the overall dominance of kinetic features.

In support of this conclusion, work from our laboratory shows that steady state flux can exert significant control over the dynamic response to perturbation (Loriaux and Hoffmann, 2013). In other words, isostatic signaling networks can exhibit very different of dynamics to the same perturbation, and only when certain protein flux parameters are constrained can a specified stimulus-response behavior be observed.

To understand why kinetic features of the regulatory network are so important in determining the cellular response to TRAIL, we constructed a minimal mechanistic model focused on the interactions of XIAP and caspase. Stability analysis of the model revealed that the de-facto positive feedback loop of caspase 3 (including the mitochondrial feedforward loop in the detailed model) functions as a strong attractor that traps transient dips of XIAP levels; a strong synthesis rate that replenishes XIAP rather than its steady state abundance ensures that XIAP levels never fall below a threshold level. As such, TRAIL responsiveness of cancer cells is governed by a kinetic buffering network motif.

In the present minimum model, the bi-stable steady state originates from a combination of the positive feedback loop and the degradation by the inhibitor. The kinetic feature controls the sensitivity of inhibitor’s abundance change to the perturbation of the time-dependent activation function and thus the degradation flux of the effector (Figure 5 and Supplementary Figure 6). We thus expect that the kinetic buffering effect revealed by the minimum model could play a role in a wide range of reaction networks that contain a similar motif. It is also related to the a buffering effect of the futile enzymatic cycle (“GK loop”) which exhibits non-linear dose responses (Xu and Gunawardena, 2012).

The present work differs from previous parameter sensitivity analyses or biomarker studies (Aldridge et al., 2006, 2011; Kallenberger et al., 2014), where initial abundances were varied. Here, model species, which are subject to synthesis and decay, were perturbed in such a way that their abundances and fluxes could be varied independently, enabling a comparison of which are more predictive. In other studies, key Hill coefficients, which are characteristic of non-linear dose responses and typically reflect more complex mechanistic control, were reported as predictive of outcome (Fey et al., 2015; Kim and Schoeberl, 2015). Here, we considered a model of mass action which does not have Hill coefficients, but then revealed that the reaction flux provides a buffering effect that can produce a non-linear dose response.

This work may help inform the development of next generation diagnostics. In cases where kinetic features are more powerful predictors of the therapeutic response than static features, then this may prompt either (i) development of novel assays or (ii) focusing on proxy or surrogate assays. Regarding the former, (i), sensitive assays for measuring kinetic features in clinical biopsies, perhaps via pulse-labeling (Doherty et al., 2009; Friedel et al., 2009; Schwanhäusser et al., 2011) may be possible. However, such diagnoses require fresh, unfixed primary human tissue samples. Regarding the latter, (ii), it may be possible to infer the values of kinetic features from other, more easily undertaken abundance measurements. For example, the rate of protein synthesis is largely determined by mRNA abundance (Ingolia et al., 2009), and the rate of degradation of a particular protein may be inferable from the abundance of the E3 ubiquitin ligase that targets it for degradation. Indeed, combination measurements such as abundances of mRNA and protein could in principle provide valuable information about protein flux. Whether such surrogate measurements are sufficient and thus more predictive than the abundance of the primary signal transducers, remains to be addressed in future studies. The present work thus may spur development of novel biomarker discovery efforts based on the recognition that it may not be the abundances but the fluxes of key regulators that control biological phenotype.

## Methods

### Workflow demonstration with a bistable model

To demonstrate the workflow of using py-substitution (Loriaux et al., 2013) to obtain the steady state solution for mass action model and study the dependence of a binary outcome on kinetic or static features, we considered a small mass-action model with bistability (Bishop and Qian, 2010). The model contains chemical reactions of an autocatalytic kinase, given by the following reactions:

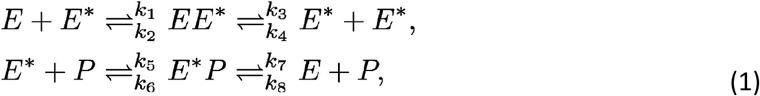

where *E* is an enzyme, *E*^∗^ is an autocatalytic kinase, *P* is a phosphatase. The model can exhibit bistability with deterministic dynamics.

Then, the mass-action model of the chemical species vector ***x*** is given by the rate equations:

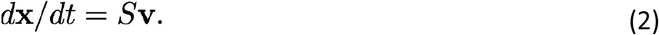

The stoichiometric matrix and the reaction fluxes are:

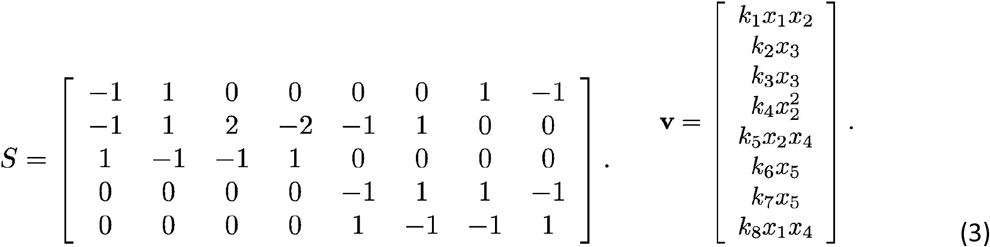

We implemented py-substitution to obtain the steady-state expression for the equation, such that the parameters can be distributed in a way to maintain the steady state. We then distributed all the independent rate constants and abundances (cf. Figure 2). The simulation of the model is able to generate bistable steady state values for the phosphatase *P*. We ran 20000 simulations, binarized the steady-state values of *P*, and implemented the regression analysis to get the correlation on the features as in the apoptosis model. The result (Supplementary Figure 1) shows that the static features have higher correlation with the phosphatase *P* binary value than the kinetic features in this model.

To investigate the model further, we added synthesis and degradation reactions to the enzyme *E* and phosphatase *P*, reflecting the reality of biochemical species (cf. Supplementary Figure 3). Then, the stoichiometric matrix and the reaction fluxes becomes:

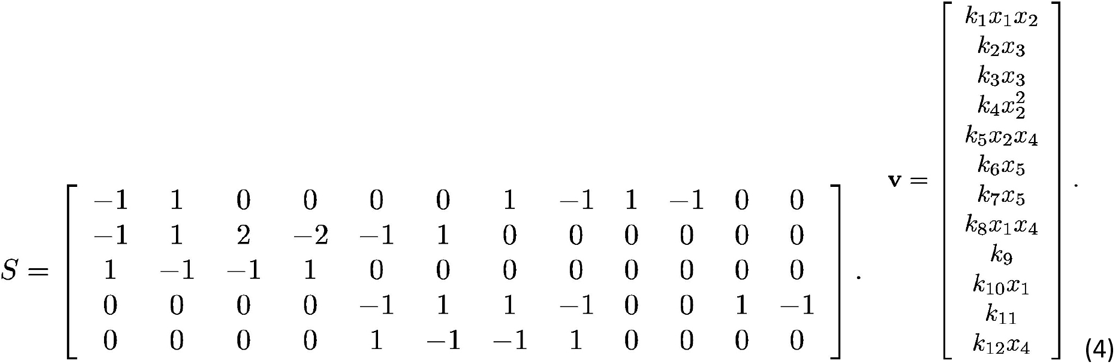

We again performed py-substitution to obtain the steady-state expression for the equation and repeated the above procedures. The result (Supplementary Figure 2) shows that the kinetic features now rank higher in the correlation with the phosphatase *P* binary value than the static features for this model. The synthesis and degradation fluxes may represent important kinetic features that correlate with the models’ binary decision.

### The mathematical model of TRAIL-induced apoptosis

To construct a bi-responsive model of TRAIL-induced apoptosis, the Albeck model (Albeck et al., 2008a) was extended to include 15 synthesis reactions, 28 degradation reactions, and deactivation reactions for the species Mito and Apaf (Supplementary Table 3). An analytical expression for the steady state was derived using Maple version 14 (Loriaux et al., 2013) and a py-substitution strategy that preserved all internal reaction kinetics and steady state species abundances as reported in the Albeck model. The 45 new reactions required 31 additional parameters, 28 of them being degradation rate constants. For these we imposed a nominal half-life of one hour, justified by the observation that signaling proteins tend to be short lived (Loriaux and Hoffmann, 2013). Several species half-lives were then manually adjusted to better fit published dynamic profiles for active caspase 8, caspase 3, and cleaved Parp. They are: the Bar-caspase 8 complex, caspase 6, cleaved Parp, the TRAIL ligand, cytoplasmic Cytochrome C, and Bar. Inactivation of Mito and Apaf were assumed to be 10 times faster than protein degradation. See Supplementary Tables 4-7 for a complete listing of parameters and their values.

Heterogeneity in the steady state was achieved by letting 14 independent steady state abundances be gamma distributed and 11 independent degradation rate constants be log normally distributed. Variance in the species abundances was set equal to one-tenth the square of the mean, i.e. the “extrinsic noise limit” observed in (Taniguchi et al., 2010). Mean abundances were taken from (Albeck et al., 2008a). For each gamma distribution, the scale parameter was calculated from the square of the mean over the variance, and the shape parameter from the variance over the mean. Degradation rate constants were assumed to have a CV of 0.368, equivalent to a variance of 1 in the log-normal distribution of protein half-lives. 40 species and 15 kinetic parameters likewise assumed a probability distribution by virtue of being constrained to steady state (Figure 2A). Only the internal reaction kinetics, as well as degradation rates of complexes and modified species, remained constant.

In using py-substitution, kinetic parameters and abundances are constrained by the steady state condition, in which fluxes are non-zero, and the conditions of detailed balance do not apply. The predictive power of kinetic features can only occur in systems without detailed balance, which is generally true for a biological system. It is also worth noting that the flux obtained from py-substitution is different from that calculated through Jacobian matrix of the mass action equation. The former represents real flux of reactive chemical species, whereas the eigenvalues of the latter are mathematical terms indicating the behavior of the solution to the linearized equation around the fixed point.

### Feature selection and regression

To sample the heterogeneous population described by our TRAIL model, values were chosen for each of the 25 independent, random parameters according to their prescribed probability density functions. These values were then used to calculate the 57 dependent parameters whose values were constrained by steady state. To simulate each sample’s response to stimulation, we instantaneously added 1000 molecules of TRAIL and numerically integrated the system to 48 hours post-stimulation using MATLAB version R2018b.

To score the outcome of the simulation, the amount of tetrameric Bax or cleaved PARP at 48 hours was recorded and fit to a mixture of two univariate Gaussians using the method of derivatives. Specifically, we first calculated a kernel density estimate of the response variable at 48 hours. The first and second derivatives of the estimate were calculated by forward finite difference and smoothed using loess locally weighted linear regression. The minima of the second derivative were taken to be the means of the two Gaussians and the distance between these and the minima of the first derivative taken to be the standard deviation. Samples that were assigned to the Gaussian with the higher mean were assigned a response variable of 1, indicating a positive response to TRAIL. Those that were assigned to the Gaussian with the lower mean were assigned a response variable of 0, indicating a negative response to TRAIL.

Once binarized, the Pearson correlation statistic was calculated between each of the 82 features and the response. Again, note that for classification, we refer to the 82 random valued parameters as features. Also, the same Pearson correlation statistic was calculated between every pair of features. The absolute values of these correlation statistics yielded a feature relevance vector *F* and a redundancy, or cross-correlation matrix *Q*, respectively. From these, a weight vector ***w*** was calculated that minimizes the quadratic program

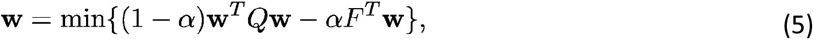

where the superscript *T* denotes the vector transpose, and the weighting factor *α* was calculated empirically by dividing the mean of *F* by the sum of *F*’s and *Q*’s mean (Mikkaichi et al., 2019).

Once the optimal vector of weights was identified, we used this as an ordering by which to incorporate features into a logistic regression model. Specifically, we modeled the log-odds ratio of the probability of responding to TRAIL versus not responding as a linear combination of the steady state features. Model accuracy was calculated as the fraction of true positive and true negatives over all predictions. ROC curves were obtained by iterating the log-odds ratio threshold over its full range of possible values (Mikkaichi et al., 2019), and at each value calculating the true and false positive rates, i.e. the ratio of true to total positives, and the ratio of false to total negatives, respectively.

### The minimal mechanistic model for analytical treatment

We constructed a model for the reactions represented in Figure 5A. The ordinary differential equations are listed here:

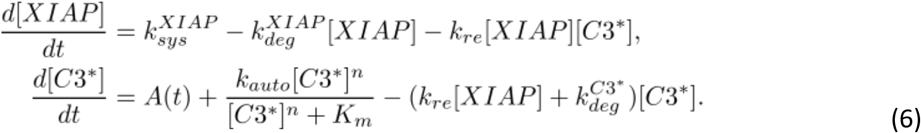

In the model, XIAP is synthesized and degraded, and consumed by the reaction catalyzed by C3*. For C3* dynamics, we adopted a Hill equation (MacArthur et al., 2009) to model the positive autoregulation of C3* by a positive feed-forward loop as indicated by (Albeck et al., 2008a). The Hill function of C3* was parameterized, such that the final steady-state value of C3* matches the extended cell death model. Without loss of generality, we chose the Hill coefficient to be 3 to model the possible cooperative effect of positive feed-forward loop. The Km value was chosen as 1e12 to ensure an auto-regulation strength with the same order of magnitude as in the original larger model.

The activation of C3* by upstream signaling was modeled by a time-dependent smooth pulse-like function. For the convenience of modeling a continuous smooth pulse function, we chose a log-normal distribution as the time-dependent activation A(t). The time scale for this pulse input is set by the reactions in the original cell death model, when the cell is stimulated by TRAIL. Thus, we parameterized the log-normal distribution such that the peak time is around 5 hours (Figure 5C).

We also used a degradation term in C3* dynamics to approximate the consumption by enzymes of other reactions, such as PARP. The degradation rate of C3* was chosen such that under the minimum overall consumption on C3*, i.e., when XIAP abundance is zero, the steady-state C3* abundance matches the simulation of the original cell death model. Other parameters had the same values as in the original model.

Since we aimed to perturb the synthesis and degradation rate of XIAP, we used py-substitution to keep the steady state abundances fixed (Loriaux and Hoffmann, 2013; Loriaux et al., 2013). Through py-substitution, we obtained an analytical solution to simultaneously vary synthesis and degradation rate of XIAP, without changing other initial conditions. This enabled studying the effects of kinetic features, without affecting abundances.

We further explored a version of the model in which XIAP is modeled enzymatically rather than stoichiometrically, that is, C3* does not degrade XIAP. To this end, we removed the last term in the first equation of XIAP dynamics. In order to be consistent with the original minimal model of the cell fate decision under the same activation flux, we changed the inhibition strength of XIAP on C3* to have a 25-fold lower numerical value. We observed that, when XIAP is treated as an enzyme, a change in its abundance, not its flux, affects the cell fate decision (Supplementary Figure 7E-F).

## Acknowledgments

The authors would like to acknowledge many helpful discussions with Peter Sorger (Harvard University) and Charles Elkan (UCSD). The study was supported by grants to AH such as the San Diego Center for Systems Biology P50 GM085763 and collaborative grants at UCLA U19AI128913, P01AI120944, and U01AI124319. PML was supported by the Department of Energy Computational Sciences Graduate Fellowship. YT was supported by a QCB Collaboratory postdoctoral Fellowship.

## Author Contributions

PML implemented the apoptosis network model, generated the data, developed the feature selection workflow, and performed the analysis. YT developed and analyzed the demonstrational example and the minimal mechanistic model. AH supervised the project. All authors wrote the paper.

## Figures and Figure Legends

**Supplementary Figure 1.**
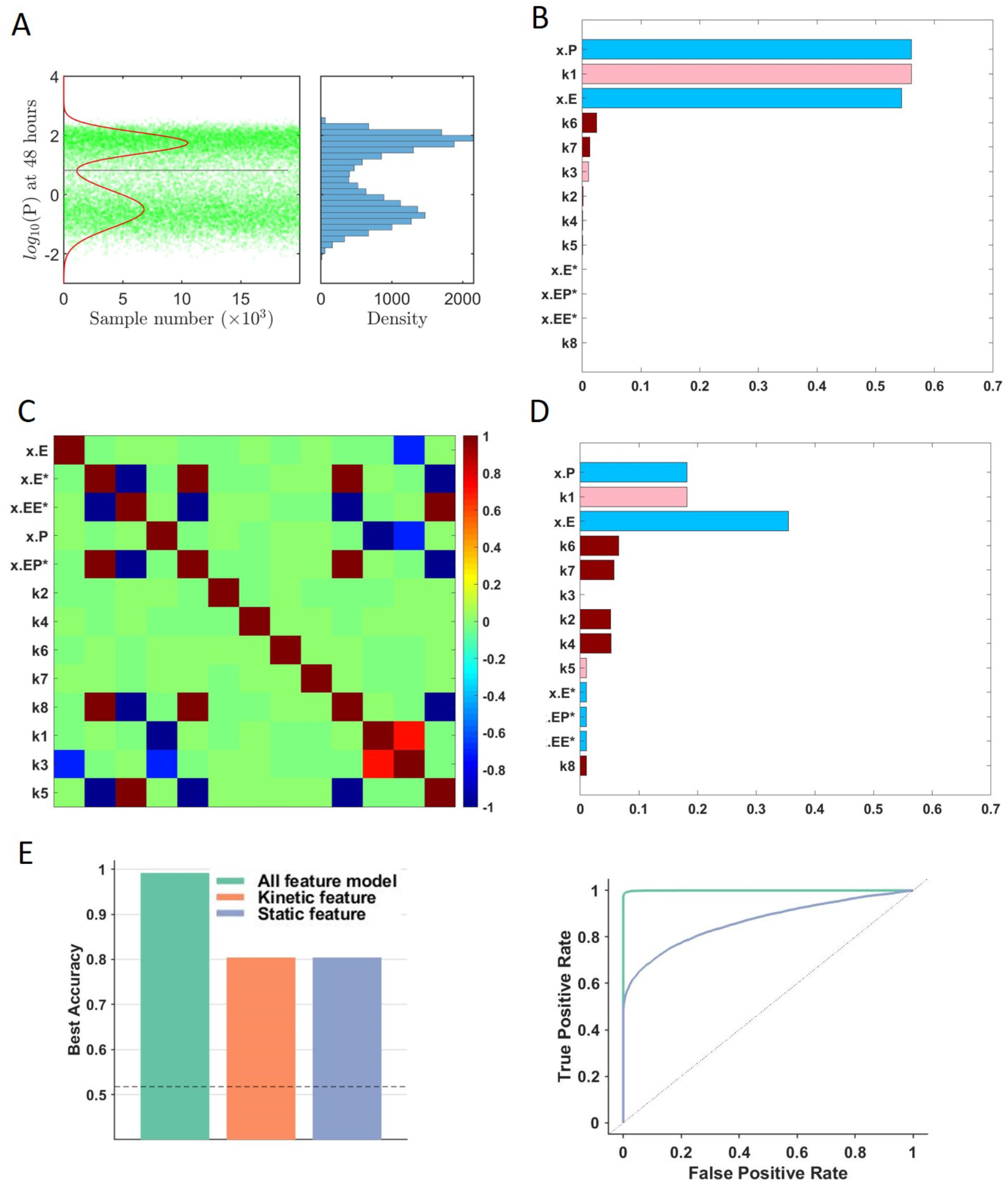
An implementation on the workflow of identifying correlated features in a bistable mass-action model. (A) The steady state values for the phosphatase *P* for a bistable mass action model (Bishop and Qian, 2010). (B) Pearson correlation coefficients between each feature and the binary response, sorted in decreasing order, as in Figure 3. The colors denote the feature types: independent species (dark blue), independent rates (dark red), dependent species (light blue) or dependent rates (light red). (C) Correlations between features. (D) QPFS weighted correlation between features and the binary response. Features are plotted in the same order (B). (E) The accuracy and the receiver operating characteristic curve of each type of the features in the logistic regression model, as in Figure 4.

**Supplementary Figure 2.**
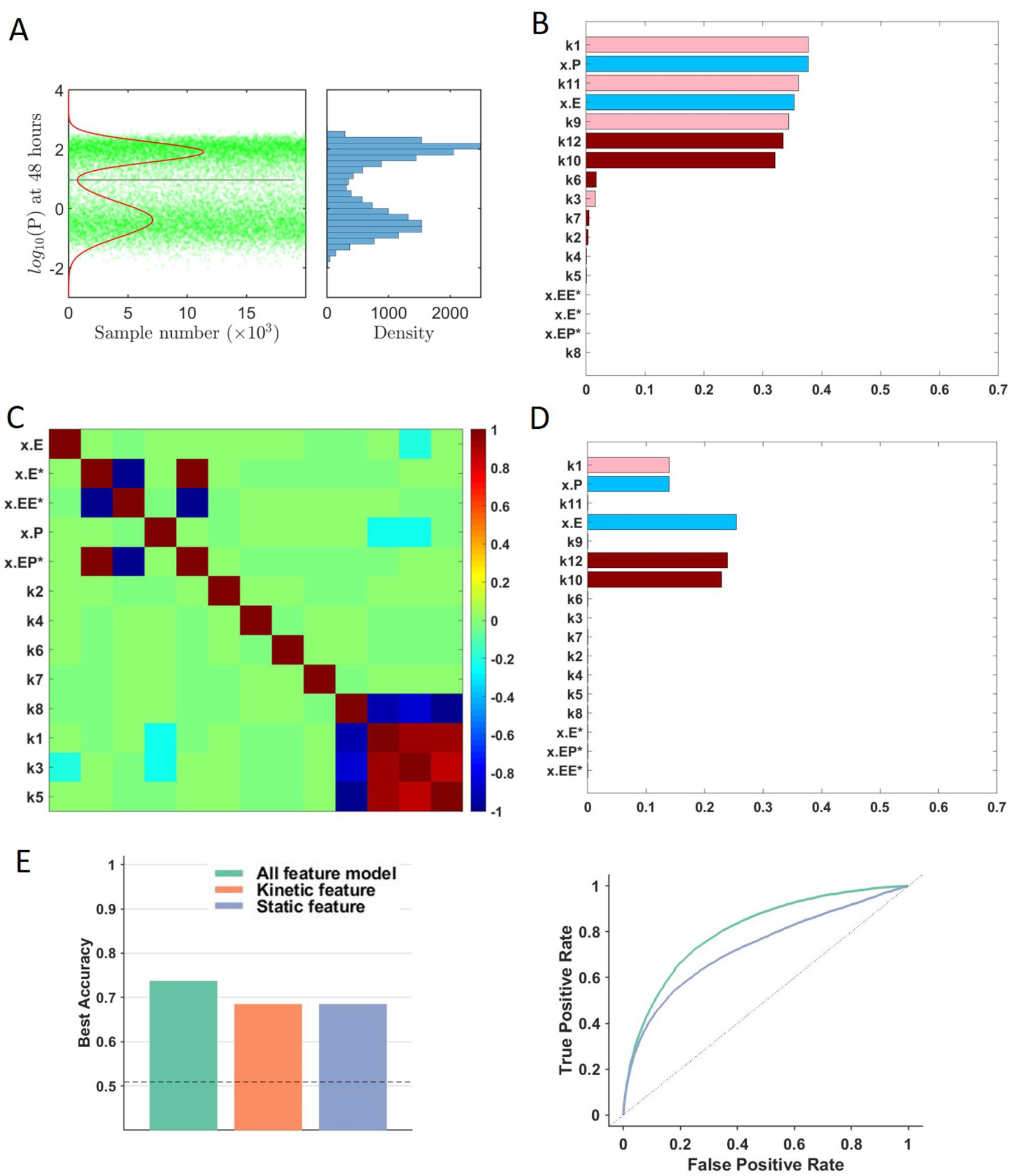
An implementation on the workflow of identifying correlated features in a bistable mass-action model with synthesis and degradation flux. (A-E) The same panels as Supplementary Figure 1 for the bistable mass-action model with synthesis and degradation flux.

**Supplementary Figure 3.**
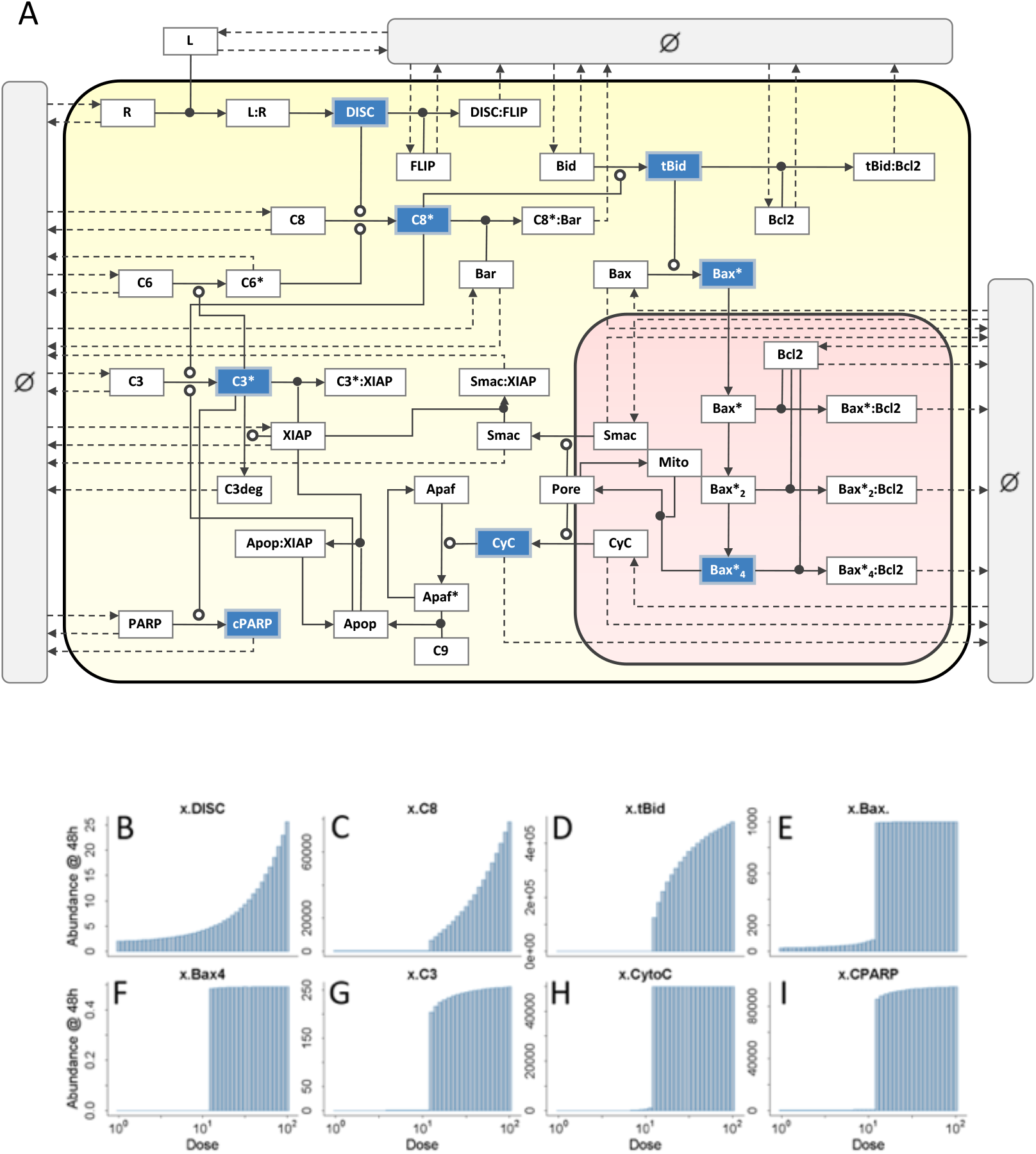
A bistable model of TRAIL-induced cell death. (A) A model of TRAIL-induced cell death that includes the original 58 species and 70 reactions introduced in (Albeck et al., 2008a), as well as 15 zero-order synthesis reactions, 28 first-order degradation reactions, and 2 first-order back-reactions resulting in deactivation of Mito and Apaf. See Supplementary Tables 1-2 for a description of all species and reactions. The 43 synthesis and degradation reactions are represented by dashed lines emanating from or terminating in a gray “null” compartment. This compartment is intended to symbolize the boundaries of the system, not the extracellular environment. Other symbols are as described in (Albeck et al., 2008a). The cytoplasm and mitochondria are shaded in yellow and rose, respectively. Species selected for dose-response analysis are shaded in blue. (B-I) Dose-response curves for the eight species in (A) shaded in blue: (B) DISC, (C) active caspase 8, (D) truncated Bid, (E) active Bax monomers in the cytoplasm (F) tetrameric Bax in the mitochondria, (G) active caspase 3, (H) Cytochrome C in the cytoplasm, and (I) cleaved PARP. Each panel depicts the absolute abundance of the species at 48 hours after stimulation with TRAIL. The dose of TRAIL ranges from 1 to 100-fold its ambient abundance.

**Supplementary Figure 4.**
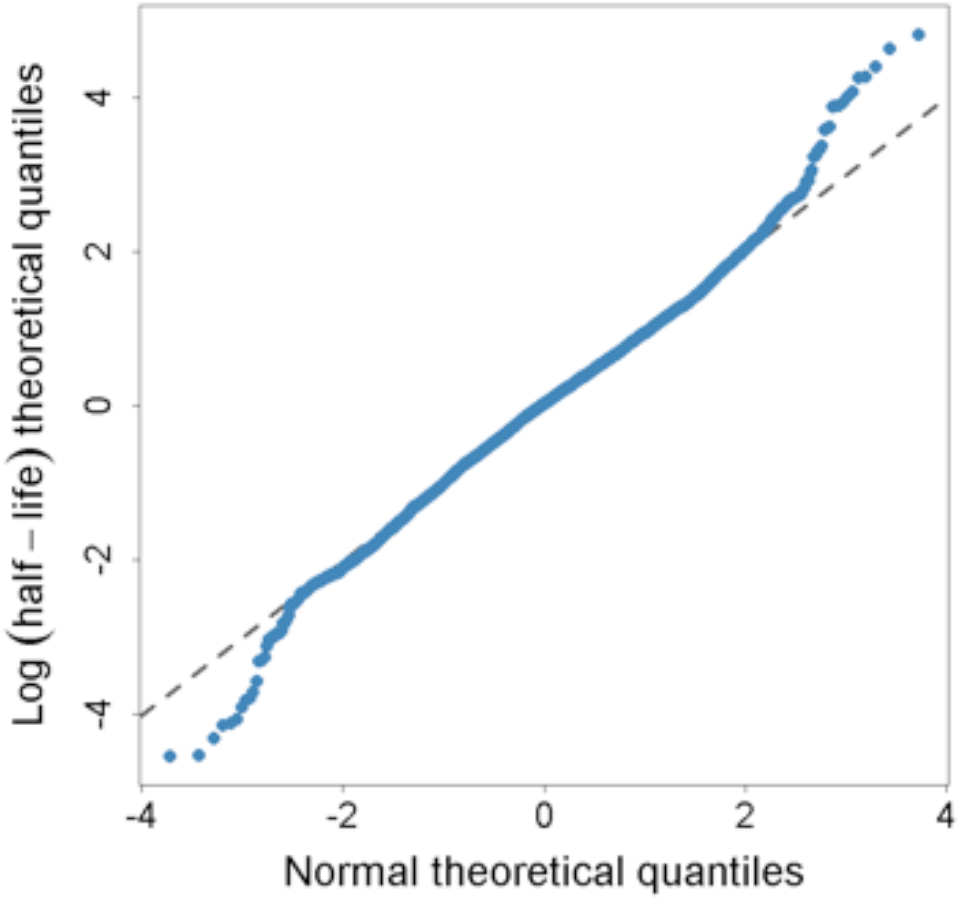
Protein half-lives are log-normally distributed. The natural log of 5028 protein half-lives, averaged over two experiments, were taken from (Schwanhäusser et al., 2011) and mean-subtracted. The quantiles of this distribution were then plotted against quantiles from a standard Normal distribution and superimposed over the line y=x (gray).

**Supplementary Figure 5.**
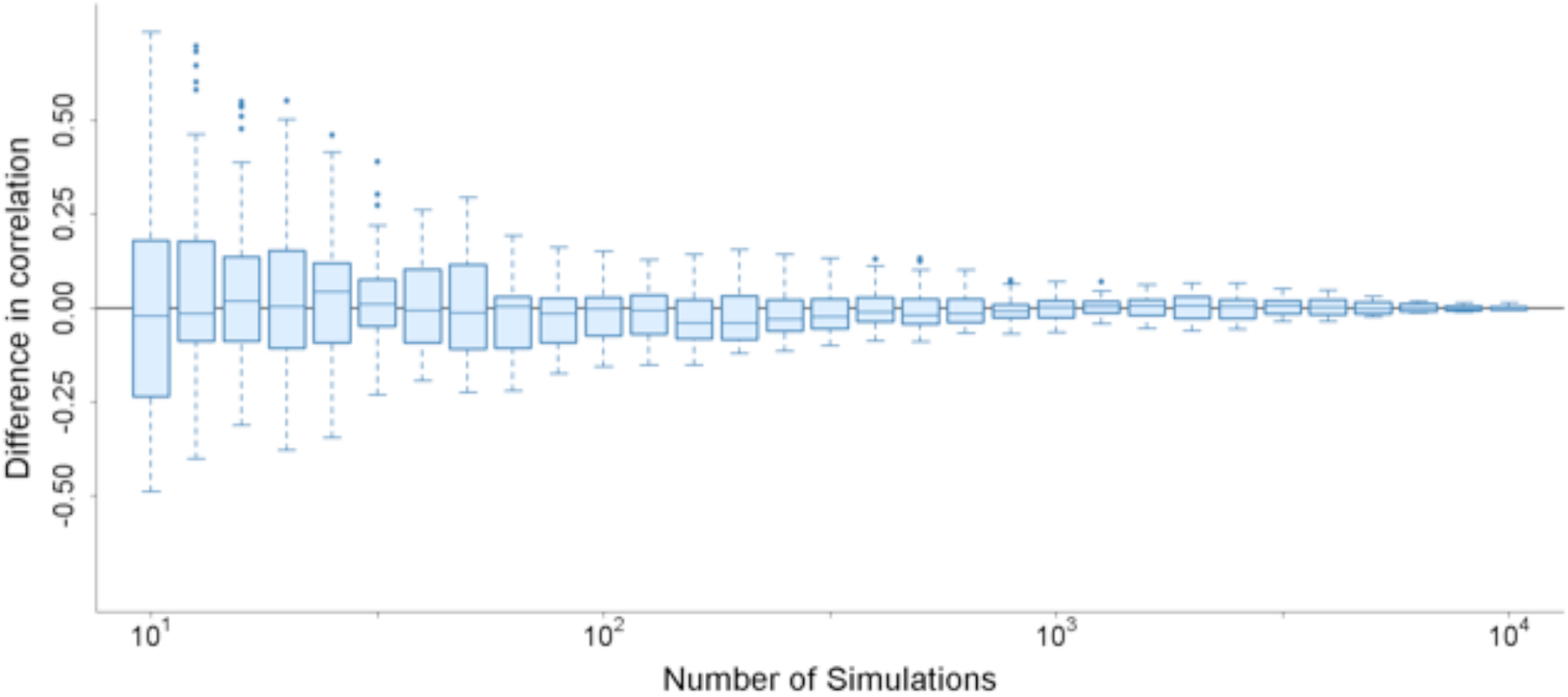
Cross-correlation between features. Pearson correlation coefficients were calculated between all pairs of features and plotted as a matrix. Hierarchical clustering was used to order the rows and columns, with clustering being performed separately for static versus kinetic features. Six clusters were identified after sorting, S1-S4, K1, and K2. We use hue to distinguish between cross-correlations within static features (blue), within kinetic features (red), and between static and kinetic features (violet). Color saturation is used to indicate the strength of the correlation, from weak (low saturation) to strong (high saturation).

**Supplementary Figure 6.**
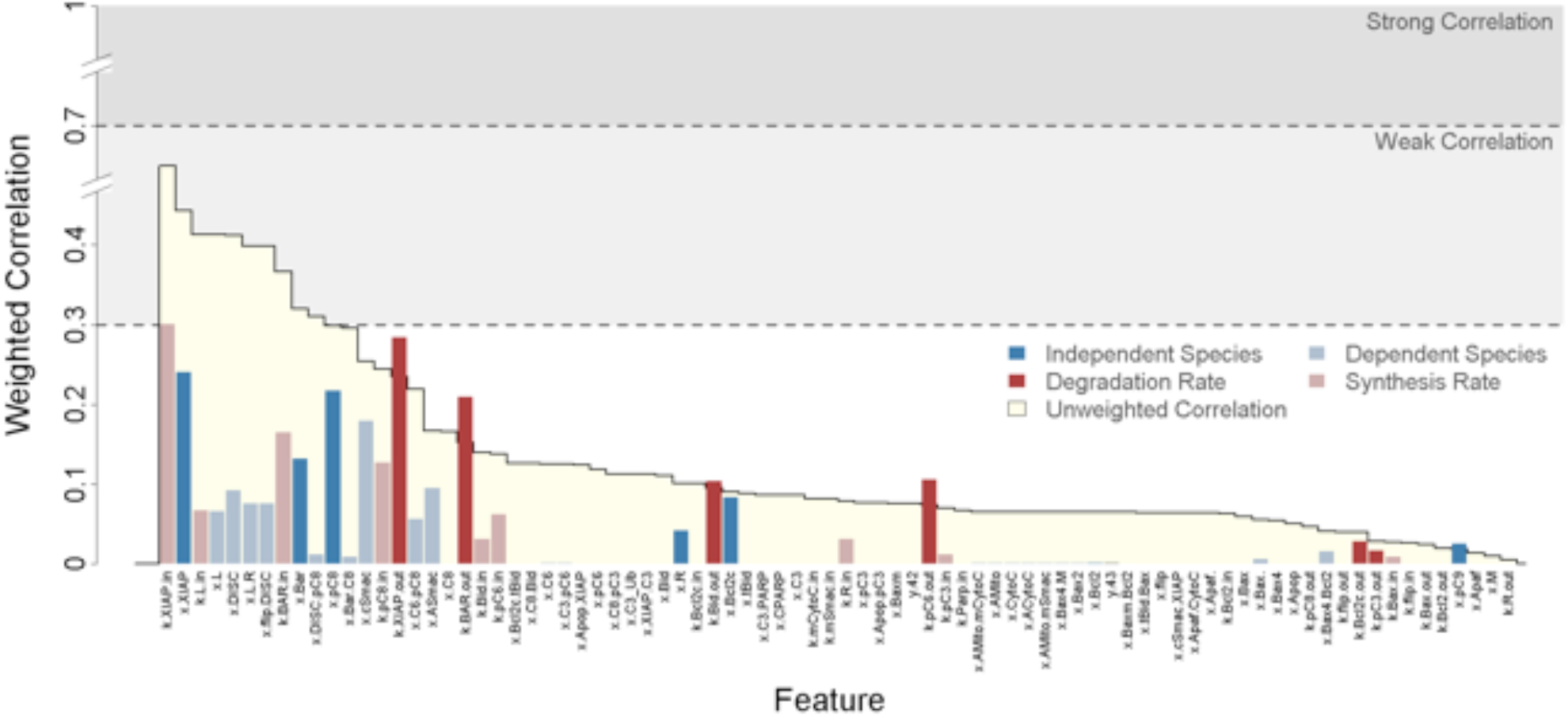
QPFS weighted correlation in the context of slow protein turnover. This figure plots QPFS-weighted correlations between each feature and the response to TRAIL in the context of 66-fold slower protein turnover than in default model. As before, independent species are dark blue, dependent species are light blue, degradation rate constants are dark red, and synthesis rates are light red.

**Supplementary Figure 7.**
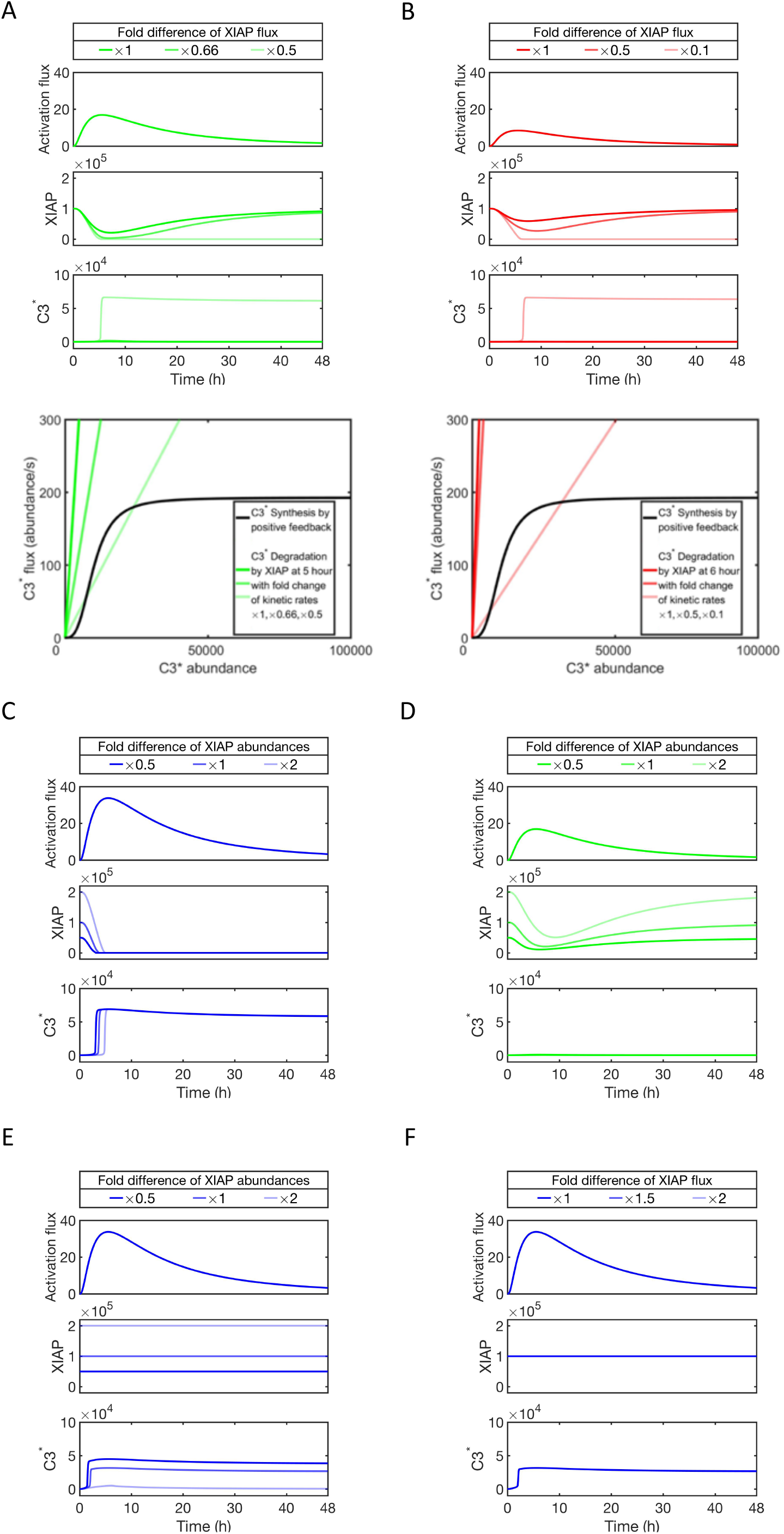
Exploring the sensitivity of kinetic or static features within the minimal model of the C3-XIAP module. (A) Slowed turnover renders cells more sensitive to sublethal TRAIL doses. Upper panel: time course of XIAP and C3* abundances under indicated of XIAP synthesis and degradation rates, given a 2-fold lower amplitude in the input activation flux than in Figure 5C. Given the XIAP synthesis and degradation rates in Figure 5B, the steady state shows high XIAP and low C3* abundances, corresponding to live cell state. Decreasing XIAP synthesis and degradation rates render cells sensitive to death. Lower panel: synthesis and degradation flux lines of C3* similar as Figure 5D. The brightness of lines denotes foldchange of XIAP synthesis and degradation rates simultaneously. With lower kinetic rates, the degradation on C3* by XIAP becomes sufficiently low, such that the steady state with higher C3* is reached. (B) The same plots as (A), with 4-fold lower amplitude of the input activation flux than in Figure 5C. Cells are sensitized to death by decreasing XIAP synthesis and degradation rates 10-fold. Lower input activation flux thus requires a more substantial decrease in XIAP kinetic rates to affect the cell fate change. (C) Steady state abundances of XIAP do not substantially affect the sensitivity of cells to lethal doses. Time course of XIAP and C3* abundances under indicated of XIAP abundances, given the input activation flux 33.77 molecules/s and 2-fold lower amplitude as Figure 5C. The varying static abundances of XIAP do not alter the cell fate decision, indicating that static feature is not predictive for the current network motif of minimum model. (D) Steady state abundances of XIAP do not substantially affect the sensitivity of cells to sublethal doses. These simulations are produced in the same conditions as (C) but with a lower activation flux. (E, F) If XIAP functioned as an enzymatic inhibitor, its abundance and not its flux, would be a key determinant of death. The plots and simulation conditions are identical to C, except XIAP is modeled as an enzyme by removing the C3*-mediated degradation reaction and correspondingly decreasing the inhibition rate of XIAP on C3*.

**Supplementary Video** Timecourses of the phase analysis at XIAP fluxes of 1x, 1.5x and 2x.

## References

Albeck, J.G., Burke, J.M., Spencer, S.L., Lauffenburger, D.A., and Sorger, P.K. (2008a). Modeling a Snap-Action, Variable-Delay Switch Controlling Extrinsic Cell Death. PLOS Biology 6, e299.

Albeck, J.G., Burke, J.M., Aldridge, B.B., Zhang, M., Lauffenburger, D.A., and Sorger, P.K. (2008b). Quantitative Analysis of Pathways Controlling Extrinsic Apoptosis in Single Cells. Molecular Cell 30, 11–25.

Alber, A.B., Paquet, E.R., Biserni, M., Naef, F., and Suter, D.M. (2018). Single Live Cell Monitoring of Protein Turnover Reveals Intercellular Variability and Cell-Cycle Dependence of Degradation Rates. Molecular Cell 71, 1079-1091.e9.

Aldridge, B.B., Haller, G., Sorger, P.K., and Lauffenburger, D.A. (2006). Direct Lyapunov exponent analysis enables parametric study of transient signalling governing cell behaviour. IEE Proceedings - Systems Biology 153, 425–432.

Aldridge, B.B., Gaudet, S., Lauffenburger, D.A., and Sorger, P.K. (2011). Lyapunov exponents and phase diagrams reveal multi-factorial control over TRAIL-induced apoptosis. Molecular Systems Biology 7, 553.

Alizadeh, A.A., and Staudt, L.M. (2000). Genomic-scale gene expression profiling of normal and malignant immune cells. Current Opinion in Immunology 12, 219–225.

Almasan, A., and Ashkenazi, A. (2003). Apo2L/TRAIL: apoptosis signaling, biology, and potential for cancer therapy. Cytokine & Growth Factor Reviews 14, 337–348.

Ashkenazi, A., and Herbst, R.S. (2008). To kill a tumor cell: the potential of proapoptotic receptor agonists. J Clin Invest 118, 1979–1990.

Auffray, C., Chen, Z., and Hood, L. (2009). Systems medicine: the future of medical genomics and healthcare. Genome Medicine 1, 2.

Bishop, L.M., and Qian, H. (2010). Stochastic Bistability and Bifurcation in a Mesoscopic Signaling System with Autocatalytic Kinase. Biophysical Journal 98, 1–11.

Boisvert, F.-M., Ahmad, Y., Gierliński, M., Charrière, F., Lamont, D., Scott, M., Barton, G., and Lamond, A.I. (2012). A Quantitative Spatial Proteomics Analysis of Proteome Turnover in Human Cells. Molecular & Cellular Proteomics 11, M111.011429.

Cambridge, S.B., Gnad, F., Nguyen, C., Bermejo, J.L., Krüger, M., and Mann, M. (2011). Systems-wide Proteomic Analysis in Mammalian Cells Reveals Conserved, Functional Protein Turnover. J. Proteome Res. 10, 5275–5284.

Chechlinska, M., Kowalewska, M., and Nowak, R. (2010). Systemic inflammation as a confounding factor in cancer biomarker discovery and validation. Nature Reviews Cancer 10, 2–3.

Diamandis, E.P. (2010). Cancer Biomarkers: Can We Turn Recent Failures into Success? J Natl Cancer Inst 102, 1462–1467.

Doherty, M.K., Hammond, D.E., Clague, M.J., Gaskell, S.J., and Beynon, R.J. (2009). Turnover of the Human Proteome: Determination of Protein Intracellular Stability by Dynamic SILAC. J. Proteome Res. 8, 104–112.

Düssmann, H., Rehm, M., Concannon, C.G., Anguissola, S., Würstle, M., Kacmar, S., Völler, P., Huber, H.J., and Prehn, J.H.M. (2010). Single-cell quantification of Bax activation and mathematical modelling suggest pore formation on minimal mitochondrial Bax accumulation. Cell Death & Differentiation 17, 278–290.

Fey, D., Halasz, M., Dreidax, D., Kennedy, S.P., Hastings, J.F., Rauch, N., Munoz, A.G., Pilkington, R., Fischer, M., Westermann, F., et al. (2015). Signaling pathway models as biomarkers: Patient-specific simulations of JNK activity predict the survival of neuroblastoma patients. Sci. Signal. 8, ra130–ra130.

Friedel, C.C., Dölken, L., Ruzsics, Z., Koszinowski, U.H., and Zimmer, R. (2009). Conserved principles of mammalian transcriptional regulation revealed by RNA half-life. Nucleic Acids Res 37, e115–e115.

Friedman, N., Cai, L., and Xie, X.S. (2006). Linking Stochastic Dynamics to Population Distribution: An Analytical Framework of Gene Expression. Phys. Rev. Lett. 97, 168302.

Gaudet, S., Spencer, S.L., Chen, W.W., and Sorger, P.K. (2012). Exploring the Contextual Sensitivity of Factors that Determine Cell-to-Cell Variability in Receptor-Mediated Apoptosis. PLOS Computational Biology 8, e1002482.

Hood, L., and Friend, S.H. (2011). Predictive, personalized, preventive, participatory (P4) cancer medicine. Nature Reviews Clinical Oncology 8, 184–187.

Ingolia, N.T., Ghaemmaghami, S., Newman, J.R.S., and Weissman, J.S. (2009). Genome-Wide Analysis in Vivo of Translation with Nucleotide Resolution Using Ribosome Profiling. Science 324, 218–223.

Kallenberger, S.M., Beaudouin, J., Claus, J., Fischer, C., Sorger, P.K., Legewie, S., and Eils, R. (2014). Intra- and Interdimeric Caspase-8 Self-Cleavage Controls Strength and Timing of CD95-Induced Apoptosis. Sci. Signal. 7, ra23–ra23.

Kaufmann, S.H., Desnoyers, S., Ottaviano, Y., Davidson, N.E., and Poirier, G.G. (1993). Specific Proteolytic Cleavage of Poly(ADP-ribose) Polymerase: An Early Marker of Chemotherapy-induced Apoptosis. Cancer Res 53, 3976–3985.

Kim, J., and Schoeberl, B. (2015). Beyond static biomarkers—The dynamic response potential of signaling networks as an alternate biomarker? Sci. Signal. 8, fs21–fs21.

King, E.L., and Altman, C. (1956). A Schematic Method of Deriving the Rate Laws for Enzyme-Catalyzed Reactions. J. Phys. Chem. 60, 1375–1378.

Lam, C.F., and Priest, D.G. (1972). Enzyme Kinetics: Systematic Generation of Valid King-Altman Patterns. Biophysical Journal 12, 248–256.

Le Novère, N., Bornstein, B., Broicher, A., Courtot, M., Donizelli, M., Dharuri, H., Li, L., Sauro, H., Schilstra, M., Shapiro, B., et al. (2006). BioModels Database: a free, centralized database of curated, published, quantitative kinetic models of biochemical and cellular systems. Nucleic Acids Res 34, D689–D691.

Lopez, C.F., Muhlich, J.L., Bachman, J.A., and Sorger, P.K. (2013). Programming biological models in Python using PySB. Molecular Systems Biology 9, 646.

Loriaux, P.M., and Hoffmann, A. (2013). A Protein Turnover Signaling Motif Controls the Stimulus-Sensitivity of Stress Response Pathways. PLOS Computational Biology 9, e1002932.

Loriaux, P.M., Tesler, G., and Hoffmann, A. (2013). Characterizing the Relationship between Steady State and Response Using Analytical Expressions for the Steady States of Mass Action Models. PLOS Computational Biology 9, e1002901.

MacArthur, B.D., Ma’ayan, A., and Lemischka, I.R. (2009). Systems biology of stem cell fate and cellular reprogramming. Nature Reviews Molecular Cell Biology 10, 672–681.

Mahalingam, D. N.A.M., Oldenhuis, C., Szegezdi, E.J. Giles, F.G.E. de Vries, E., de Jong, S., and T.Nawrocki, S. (2011). Targeting Trail Towards the Clinic.

Mandal, M., Maggirwar, S.B., Sharma, N., Kaufmann, S.H., Sun, S.-C., and Kumar, R. (1996). Bcl-2 Prevents CD95 (Fas/APO-1)-induced Degradation of Lamin B and Poly(ADP-ribose) Polymerase and Restores the NF-κB Signaling Pathway. J. Biol. Chem. 271, 30354–30359.

Mikkaichi, T., Yeaman, M.R., Hoffmann, A., and Group, M.S.I. (2019). Identifying determinants of persistent MRSA bacteremia using mathematical modeling. PLOS Computational Biology 15, e1007087.

Piccart-Gebhart, M.J., Procter, M., Leyland-Jones, B., Goldhirsch, A., Untch, M., Smith, I., Gianni, L., Baselga, J., Bell, R., Jackisch, C., et al. (2005). Trastuzumab after Adjuvant Chemotherapy in HER2-Positive Breast Cancer. New England Journal of Medicine 353, 1659–1672.

Porter, A.G., and Jänicke, R.U. (1999). Emerging roles of caspase-3 in apoptosis. Cell Death and Differentiation 6, 99–104.

Rifai, N., Gillette, M.A., and Carr, S.A. (2006). Protein biomarker discovery and validation: the long and uncertain path to clinical utility. Nature Biotechnology 24, 971–983.

Rodriguez-Lujan, I., Huerta, R., Elkan, C., and Cruz, C.S. (2010). Quadratic Programming Feature Selection. Journal of Machine Learning Research 11, 1491–1516.

Romond, E.H., Perez, E.A., Bryant, J., Suman, V.J., Geyer, C.E., Davidson, N.E., Tan-Chiu, E., Martino, S., Paik, S., Kaufman, P.A., et al. (2005). Trastuzumab plus Adjuvant Chemotherapy for Operable HER2-Positive Breast Cancer. New England Journal of Medicine 353, 1673–1684.

Rosenwald, A., and Staudt, L.M. (2003). Gene Expression Profiling of Diffuse Large B-Cell Lymphoma. Leukemia & Lymphoma 44, S41–S47.

Schwanhäusser, B., Busse, D., Li, N., Dittmar, G., Schuchhardt, J., Wolf, J., Chen, W., and Selbach, M. (2011). Global quantification of mammalian gene expression control. Nature 473, 337–342.

Sheridan, C., and Martin, S.J. (2008). Commitment in apoptosis: slightly dead but mostly alive. Trends in Cell Biology 18, 353–357.

Spencer, S.L., Gaudet, S., Albeck, J.G., Burke, J.M., and Sorger, P.K. (2009). Non-genetic origins of cell-to-cell variability in TRAIL-induced apoptosis. Nature 459, 428–432.

Sreenath, S.N., Cho, K.-H., and Wellstead, P. (2008). Modelling the dynamics of signalling pathways. Essays In Biochemistry 45, 1–28.

Taniguchi, Y., Choi, P.J., Li, G.-W., Chen, H., Babu, M., Hearn, J., Emili, A., and Xie, X.S. (2010). Quantifying E. coli Proteome and Transcriptome with Single-Molecule Sensitivity in Single Cells. Science 329, 533–538.

Volkenstein, M.V., and Goldstein, B.N. (1966). A new method for solving the problems of the stationary kinetics of enzymological reactions. Biochimica et Biophysica Acta (BBA) - General Subjects 115, 471–477.

Wagner, K.W., Punnoose, E.A., Januario, T., Lawrence, D.A., Pitti, R.M., Lancaster, K., Lee, D., von Goetz, M., Yee, S.F., Totpal, K., et al. (2007). Death-receptor O-glycosylation controls tumor-cell sensitivity to the proapoptotic ligand Apo2L/TRAIL. Nature Medicine 13, 1070–1077.

Wang, S., and El-Deiry, W.S. (2003). TRAIL and apoptosis induction by TNF-family death receptors. Oncogene 22, 8628–8633.

Wiley, S.R., Schooley, K., Smolak, P.J., Din, W.S., Huang, C.-P., Nicholl, J.K., Sutherland, G.R., Smith, T.D., Rauch, C., Smith, C.A., et al. (1995). Identification and characterization of a new member of the TNF family that induces apoptosis. Immunity 3, 673–682.

Woo, M., Hakem, R., Soengas, M.S., Duncan, G.S., Shahinian, A., Kägi, D., Hakem, A., McCurrach, M., Khoo, W., Kaufman, S.A., et al. (1998). Essential contribution of caspase 3/CPP32 to apoptosis and its associated nuclear changes. Genes Dev. 12, 806–819.

Xu, Y., and Gunawardena, J. (2012). Realistic enzymology for post-translational modification: Zero-order ultrasensitivity revisited. Journal of Theoretical Biology 311, 139–152.

